# SIA: Selection Inference Using the Ancestral Recombination Graph

**DOI:** 10.1101/2021.06.22.449427

**Authors:** Hussein A. Hejase, Ziyi Mo, Leonardo Campagna, Adam Siepel

## Abstract

Detecting signals of selection from genomic data is a central problem in population genetics. Coupling the rich information in the ancestral recombination graph (ARG) with a powerful and scalable deep learning framework, we developed a novel method to detect and quantify positive selection: **S**election **I**nference using the **A**ncestral recombination graph (SIA). Built on a Long Short-Term Memory (LSTM) architecture, a particular type of a Recurrent Neural Network (RNN), SIA can be trained to explicitly infer a full range of selection coefficients, as well as the allele frequency trajectory and time of selection onset. We benchmarked SIA extensively on simulations under a European human demographic model, and found that it performs as well or better as some of the best available methods, including state-of-the-art machine-learning and ARG-based methods. In addition, we used SIA to estimate selection coefficients at several loci associated with human phenotypes of interest. SIA detected novel signals of selection particular to the European (CEU) population at the *MC1R* and *ABCC11* loci. In addition, it recapitulated signals of selection at the *LCT* locus and several pigmentation-related genes. Finally, we reanalyzed polymorphism data of a collection of recently radiated southern capuchino seedeater taxa in the genus *Sporophila* to quantify the strength of selection and improved the power of our previous methods to detect partial soft sweeps. Overall, SIA uses deep learning to leverage the ARG and thereby provides new insight into how selective sweeps shape genomic diversity.

## Introduction

The ability to accurately detect and quantify the influence of selection from genomic sequence data enables a wide variety of insights, ranging from understanding historical evolutionary events to characterizing the functional and disease relevance of observed or potential genetic variants. Adaptive evolution is driven by increases in frequency of alleles that enhance reproductive fitness. In addition, alleles experiencing such positive selection often provide insights into the functional or mechanistic basis of phenotypes of interest. Examples of genetic determinants of important phenotypic traits under selection in human populations include a family of mutations in the hemoglobin-β cluster, which confer resistance to malaria and are at high frequencies in many populations [1,2], loci controlling growth factor signaling pathways that contribute to short stature in Western Central African hunter-gatherer populations [3,4], as well as mutations in several genes involved in immunity, hair follicle development, and skin pigmentation [5] (reviewed in refs. [6–9]).

Population genetic methods predominantly identify positive selection through the detection of selective sweeps. As the frequency of an advantageous allele increases, linked variants in the vicinity can “hitchhike” to high frequency, leading to local reductions in genetic diversity. Previous approaches to detecting selective sweeps (such as traditional summary statistics [10], approximate likelihood and Approximate Bayesian Computation (ABC) methods [11], or supervised machine learning (ML) methods [12,13]) exploit the effect of genetic hitchhiking on the spatial haplotype structure and site frequency spectrum (SFS). Summary statistics have the advantage of being fast and easy to compute, but may confound the effects of selection on genetic diversity with the effects of complex demographic histories including bottlenecks, population expansions and structured populations. Besides, they cannot easily be used to estimate the value of the selection coefficient. Approximate likelihood and ABC methods, on the other hand, can provide an estimate of the strength of selection by aggregating multiple summary statistics [11], but can be prohibitively computationally expensive when applied at a large scale. ML methods for inferring selection can be more scalable, and can capture complex nonlinear relationships among features. With the exception of a handful of recently developed methods that operate on the multiple sequence alignment itself [14,15], however, the majority of ML approaches to selection inference solely make use of traditional summary statistics as features for prediction. In short, previous methods (including ABC and most ML methods) predominantly rely on low-dimensional summary statistics, which, even in combination, capture only a small portion of the information in the sequence data.

Recently, a new generation of inference methods have made it possible to go beyond summary statistics and estimate or sample a full ancestral recombination graph (ARG) [16–18] for a collection of sequences of interest. The ARG is a complex data structure that summarizes the shared evolutionary history and recombination events that have occurred in a collection of DNA sequences, and therefore contains highly informative features that can potentially be leveraged to make accurate inferences about selection. The ARG representation is interchangeable with a sequence of local genealogies along the genome and the recombination events that transform each genealogy to the next. The influence of selection on each allele can be characterized from the ARG, based on departures from the patterns of coalescence and recombination expected under neutrality as reflected in the local genealogies. Traditional ARG inference methods [19–23] were restricted in accuracy and scalability, limiting the practical application of ARGs. Recent advances [24], however, have enabled scalable yet statistically rigorous genome-wide ARG inference with dozens of genomes. Moreover, methods such as Relate [25] and tsinfer [26] have further dramatically improved the scalability of ARG inference to accommodate thousands or even hundreds of thousands of genomes. The latest progress in genealogical inference has paved the way for ARG-based methods to address many different questions in population genetics [24–27].

One natural way to exploit the richness of the ARG representation in inference of selection would be to extract features from inferred ARGs and feed them into a modern supervised machine-learning framework. Deep-learning methods, in particular, have recently achieved unprecedented success on a variety of challenging problems, including image recognition, machine translation, and game-play [28]. Deep learning is also highly flexible, providing many opportunities for the design of novel model architectures motivated by biological knowledge. An ARG-guided deep-learning model could potentially provide new insight into how natural selection impacts the human genome, human diseases and other phenotypes, and human evolution.

With these goals in mind, we developed a new method, called SIA (**S**election **I**nference using the **A**ncestral recombination graph), that uses a Recurrent Neural Network (RNN) [29,30] to infer the selection coefficient and allele frequency trajectory of a variant that maps to a gene tree embedded in an ARG. Rather than relying on traditional sequence-based summary statistics, SIA makes use of features based on the local genealogies extracted from the ARG. Based on these local topological features, SIA learns to infer the selection coefficient and allele frequency trajectory of a beneficial variant (see **Figure 1**). As described below, SIA performs well on benchmarks and is reasonably robust to model misspecification. Applying SIA to data from the 1000 Genomes Northern and Western European (CEU) population, we identified new and known loci under positive selection that are associated with a variety of phenotypes and estimated selection coefficients at these loci. In addition, using SIA, we built on our previous work [31] on a bird species-complex in the genus *Sporophila* by elucidating the strength and targets of selection at specific loci tied to a collection of rapid speciation events. Overall, SIA is the first method that couples ARG-based features with a machine-learning approach for population genetic inference.

**Figure 1:**
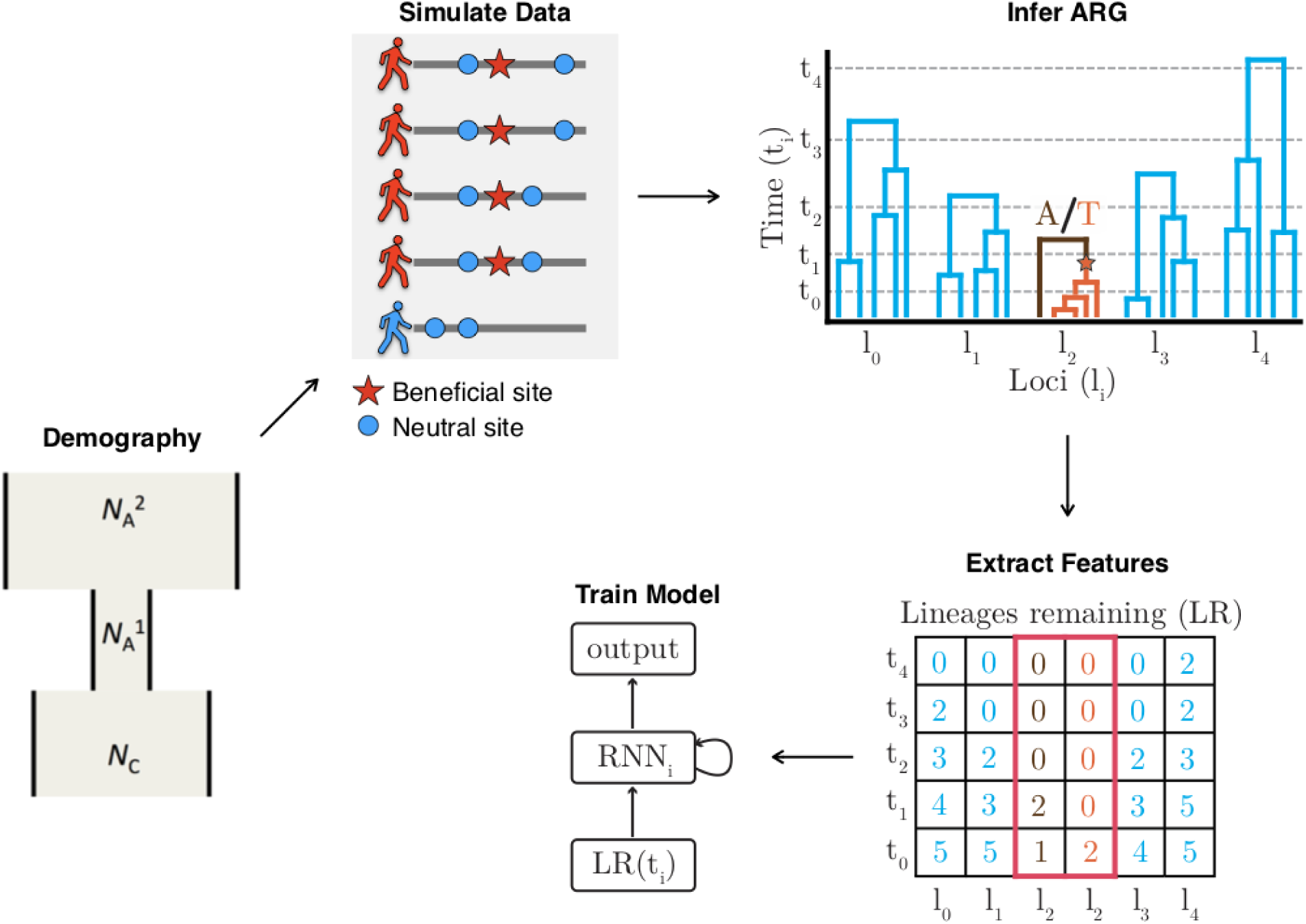
A high-level framework for automating the detection of selective sweeps. We first estimate the demographic history for the population of interest, then based on the estimated demographic history, we simulate neutral regions and sweeps using the discoal simulator [52]. We proceed with ARG inference and then extract ARG-level statistics from each simulated region. The ARG-level statistics were used as features for a deep-learning Recurrent Neural Network (RNN) model. Finally, the learned model was applied to the empirical data to infer sweeps, selection coefficients, and allele-frequency trajectories.

## Results

### Methodological overview

SIA is based on an RNN that is trained to predict selection at a genomic site from genealogical features at that site of interest and nearby sites (see **Methods** for detailed descriptions, see **Figure 1** for a conceptual overview of SIA, and **Figure S1** for an illustration of ARG features and the RNN architecture). Based on the demography of a particular population of interest, training data including genomic regions under various strengths of selection are simulated. The ARG is then inferred from each simulated data set. ARG-level statistics are extracted at the site under selection (or a neutral site) as features to be used as input to the deep-learning model. Specifically, we use lineage counts at a set of discrete time points as a fixed-dimension encoding of a genealogy. The encoding of the genealogy at the focal site as well as similar encodings of flanking genealogies constitute the feature vector for that site. SIA uses a Long Short-Term Memory (LSTM) architecture, designed specifically to handle the temporal nature of the feature set. The LSTM unrolls temporally such that the lineage counts at each time point are fed to the network iteratively. Finally, the model trained on simulations is applied to ARGs inferred from empirical data to identify sweeps, infer selection coefficients, and allele-frequency trajectories.

### Classification of sweeps

We first compared SIA with several existing methods, including the Tajima’s D [10] and H1 [32] summary statistics, iHS [33], a genealogy-based statistic [25] and a summary-statistic-based machine-learning method [12,13] (see **Methods**), in the classification task of distinguishing hard sweeps from neutrally evolving regions. Our performance comparison was conducted across 16 combinations of selection coefficients and segregating allele frequencies such that the beneficial site was subjected to selection ranging from weak to strong, resulting in low to high derived allele frequencies (DAFs). Since *a priori* we expected sweep sites with lower selection coefficients and lower DAFs to be harder to detect, we performed a stratified analysis of SIA’s performance by selection coefficient and DAF. **Figure 2** reports the Receiver Operating Characteristic (ROC) curves using simulations based on the CEU demographic model [34] where inferred genealogies were used as input to SIA to account for gene tree uncertainty. As expected, all methods tended to perform better in a regime with higher selection coefficients and DAFs, as indicated by increasing values of the area under the ROC curve (AUROC) statistic from left to right (increasing selection) and from top to bottom (increasing DAF). SIA outperformed the other methods across model conditions, with a more pronounced performance advantage for sites under weaker selection and segregating at lower DAFs (**Figure 2**). For each given selection coefficient, the AUROC of the Relate tree statistic (shown in red in **Figure 2**), which measures how unlikely it is that the observed expansion of the derived lineages is purely due to genetic drift, did not substantially improve as the DAF increased. Alleles at higher frequency tend to be older and subjected to drift over longer periods, which may lead to reduced power for Relate to distinguish lineage expansion under selection from the neutral expectation. Consequently, while the ARG-based methods SIA and Relate both outperformed other methods at low DAFs, SIA was alone in maintaining this advantage at higher DAFs.

**Figure 2:**
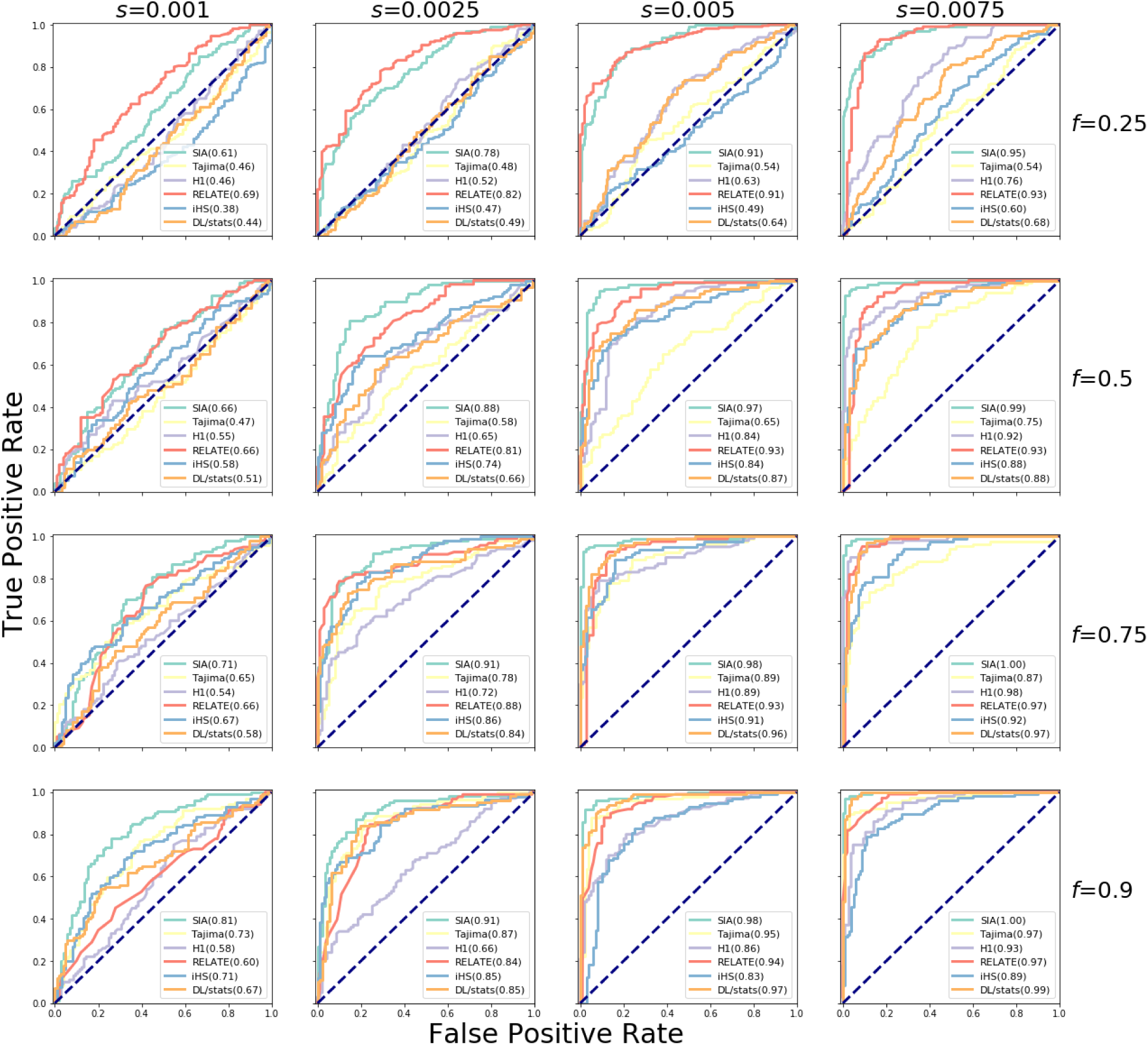
Classification performance of SIA and other methods on simulated data. Sequence data were simulated under a variety of selection regimes (*s*, shown horizontally) and derived allele frequencies (DAFs) for the beneficial mutation under selection (*f*, shown vertically) (see **Methods** for more details). The prediction task distinguished neutral regions and sweeps. The methods were tested on a set of 200 regions per panel (100 per class), and the receiver operating characteristic (ROC) curve records the true positive rate (TPR) as a function of the false positive rate (FPR). The curve is obtained by varying the prediction threshold from 0 to 1 and recording for each threshold the number of regions correctly assigned (TPs) or misassigned (FPs) as positives (with prediction probability above the threshold). The performance of each method was evaluated based on the area under its ROC curve, or AUROC. We report each method’s AUROC as an average across 200 replicate datasets for each model condition. Note that inferred genealogies were used as input to SIA.

In addition, we validated the ability of SIA to classify genomic regions with additional test sets simulated under a demographic model for southern capuchinos, a group of songbirds in which we previously identified and characterized many examples of sweeps [31], finding a predominance of “soft” rather than “hard” sweeps (meaning that they tend to be based on standing genetic variation rather than new mutations; see **Methods**). **Figure S2** reports the ROC curves for the task of distinguishing partial soft sweeps from neutral regions. Despite soft sweeps being harder to detect, the classifier achieved good performance in the moderate-to-strong selection regimes (*s* = 0.005 and *s* = 0.0075) where the accuracy ranged between 82% and 96%, a substantial improvement over the previous accuracy of 56% [31]. SIA performed particularly well in identifying partial soft sweeps when the site under selection was at a high segregating frequency. For example, at segregating frequencies of 0.75 and 0.9, the performance of SIA ranged between 80% and 96% across a variety of selection regimes (*s* = 0.0025, 0.005, and 0.0075). The performance of SIA degraded somewhat for weak selection (*s* = 0.001) with an accuracy ranging between 63% and 74%.

### Selection coefficient inference using true gene trees

We assessed the performance of SIA in correctly predicting the selection coefficient and compared it to CLUES [35]. Like SIA, CLUES uses local genealogies based on the ARG to infer a selection coefficient. However, CLUES calculates the likelihood of the genealogy analytically using a hidden Markov model (HMM), and does not rely on simulated training data. In addition, CLUES uses a single genealogy at the focal site, whereas SIA additionally considers flanking trees.

We began by supplying both methods with true genealogies, in order to later disentangle the error deriving from the ARG inference step from other sources of error (see **Discussion**). We found that SIA identified regions under neutrality with approximately no bias (median inferred *s* = 7.5e-05; **Figure 3**). Similarly, SIA correctly inferred the selection coefficient for regions under moderate to strong selection (*s* ∈ {0.0025, 0.005, 0.0075, 0.01}) with the median inferred *s* deviated from the true *s* by at most 3%. On the other hand, SIA somewhat underestimated the selection coefficient (median inferred *s* = 0.00037) for the weak selection regime (true *s* = 0.001), likely owing to limits in the training set within that selection regime (see **Discussion**). We further binned the results by segregating frequency and selection coefficient and found that, in general, the variance in estimates of *s* for SIA (as well as CLUES) tended to decrease as the segregating frequency of the beneficial allele increased (**Figure S3**).

**Figure 3:**
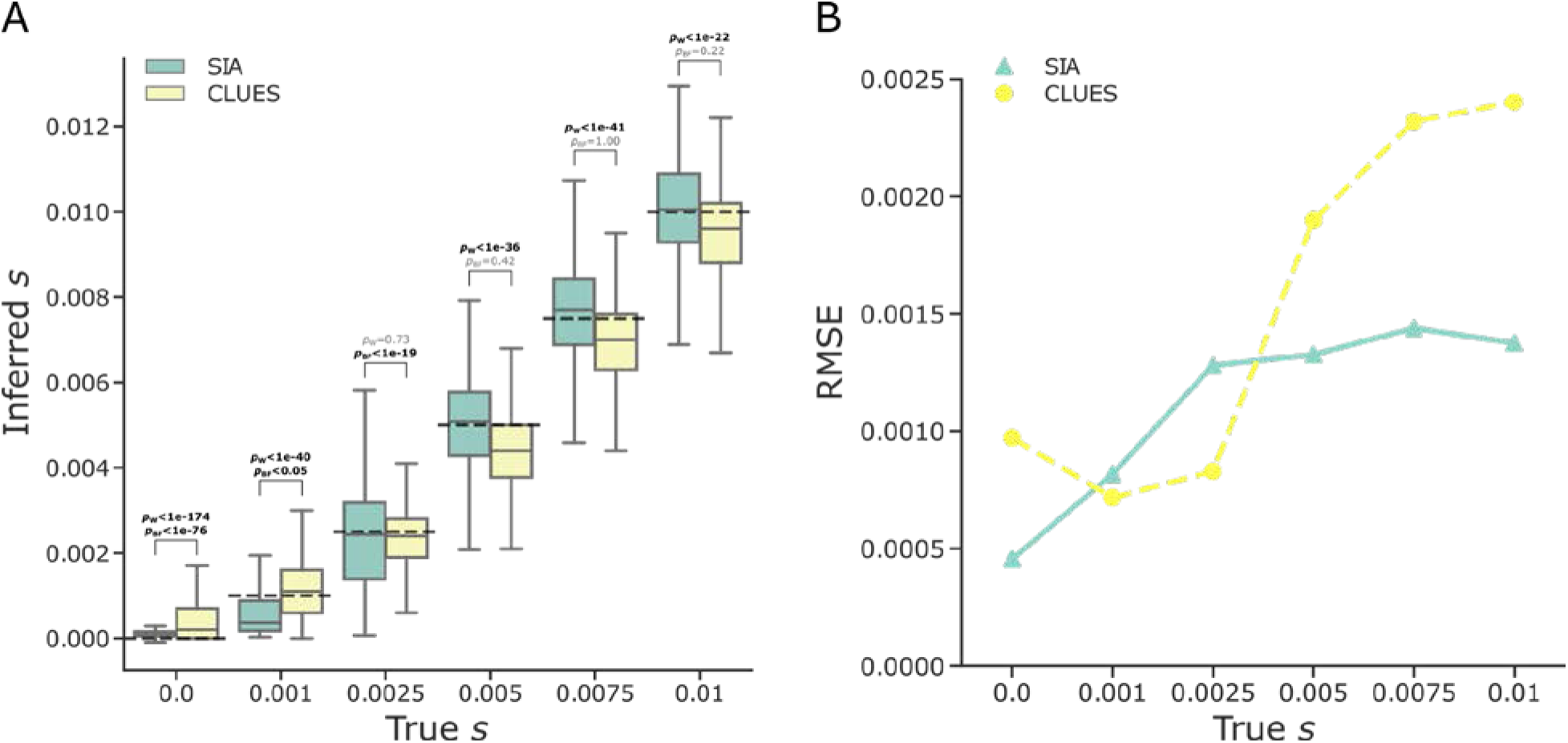
Predictions of selection coefficients for simulated regions using SIA and CLUES based on true genealogies. **(A)** The distribution of inferred selection coefficients for each method under each model condition are reported using a box plot. The box plot for each method reports these five statistics (from bottom to top): minimum, first quartile, median, third quartile, and maximum. The *y*-axis shows the inferred selection coefficient while the *x*-axis shows the true selection coefficient. The dashed-black line indicates the true selection coefficient for each model condition. The simulations are based on the CEU demographic model and true genealogies were used as input to both methods. Each model condition (i.e. box plot) represents a set of 400 independent simulations. The mean ranks and variances of the distributions of inferred *s* were compared using the Wilcoxon signed-rank test (*p*_W_) and the Brown-Forsythe test (*p*_BF_), respectively. **(B)** The root mean square error (RMSE) for each method under each model condition evaluated on 400 independent simulations.

CLUES performed roughly similarly to SIA in this experiment, but tended to slightly overestimate *s* for the neutral regions (i.e., true *s* = 0) and underestimate *s* for the moderate to high selection regimes (i.e., true *s* = 0.005, 0.0075, and 0.01). Under these conditions, SIA’s median predictions of *s* were noticeably closer to the true values (**Figure 3A**). At the same time, CLUES performed slightly better than SIA in weak selection regimes (i.e., true *s* = 0.001 and 0.0025) (**Figure 3**). Overall, SIA (RMSE = 9.52e-4) achieved a lower error in estimating *s* than CLUES (RMSE = 1.44e-3), when true genealogies were used as input to both methods (Wilcoxon signed-rank test for difference in mean of squared error, *p* = 1.25e-42). This finding potentially reflects the benefit of linkage information utilized by SIA through the additional flanking genealogies (see **Discussion**).

### Selection coefficient inference using inferred gene trees

To account for gene-tree uncertainty, we next used ARGs inferred with Relate, which is scalable to the size of the training dataset for SIA (see **Methods**), as input to SIA and CLUES and compared their performance on CEU simulations. Furthermore, we compared both methods to a supervised machine learning method, ImaGene (see **Figure S20**), that operates directly on an image of the alignment itself. ImaGene does not require gene trees as input and instead uses a Convolutional Neural Network (CNN) to perform dimensionality reduction of the sequence alignment, allowing for accurate and efficient classification and regression.

Overall, we found that SIA and ImaGene outperformed CLUES in these experiments (**Figure 4**). CLUES tended to underestimate selection coefficients for the moderate-to-strong selection regimes, to a greater extent compared to the case where true genealogies were used for inference (**Figures 3A & 4A**). This decrease in performance of CLUES evidently derives from error at the ARG reconstruction step. SIA, on the other hand, appeared to be more robust to the same ARG reconstruction error. ImaGene performed remarkably similarly to SIA, given that it relies solely on the sequence alignment. SIA exhibited lower error at neutral sites and sites with low-to-moderate values of *s*, whereas ImaGene prevailed at sites under strong selection (**Figure 4B**).

**Figure 4:**
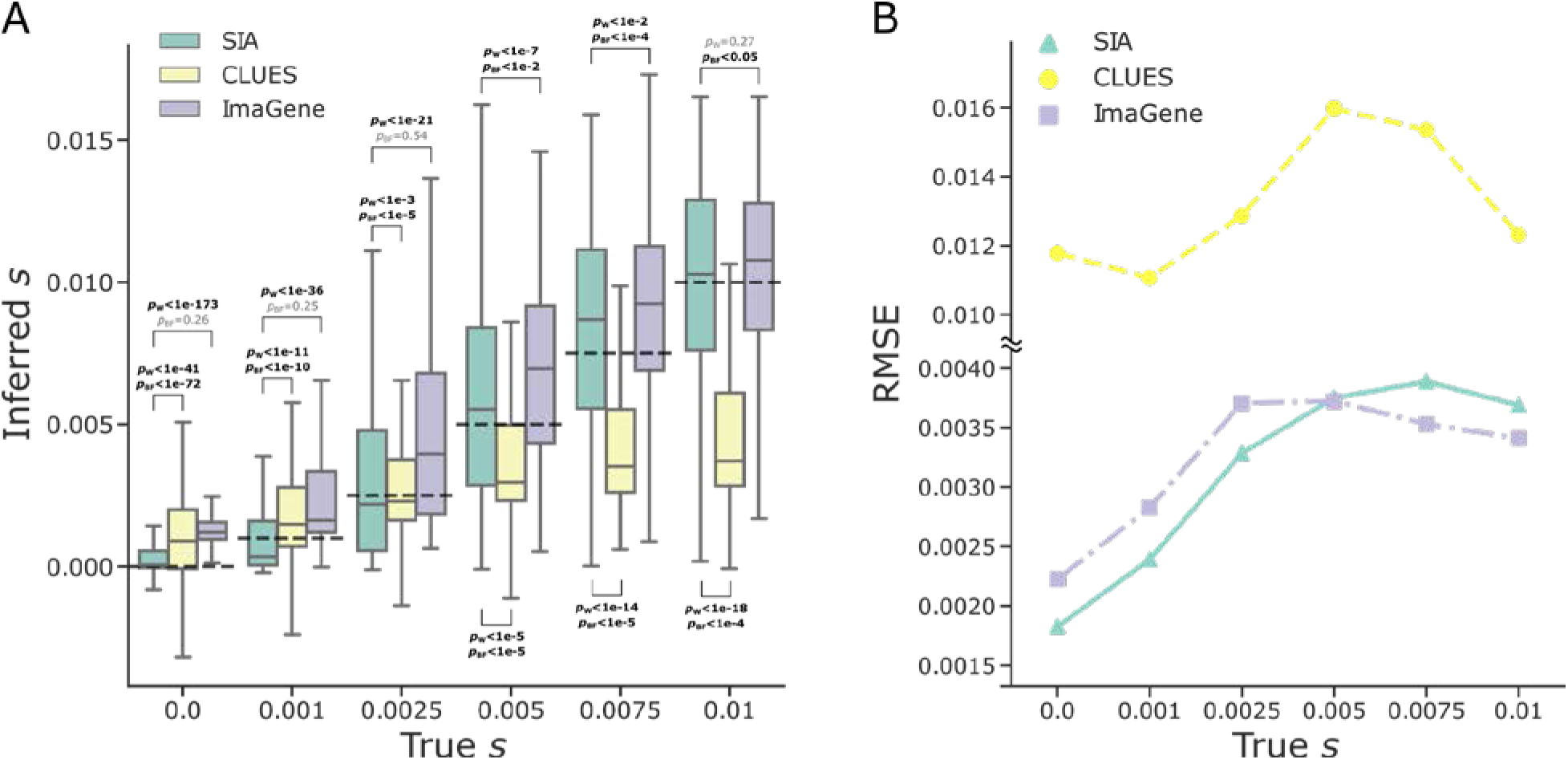
Predictions of selection coefficient on simulated regions using SIA and CLUES based on inferred genealogies, and ImaGene. **(A)** The distribution of inferred selection coefficients and **(B)** root mean square error (RMSE) for each method under each model condition. The simulations are based on the CEU demographic model where inferred genealogies were used as input to SIA and CLUES, whereas sequence alignments were used as input to ImaGene. Figure layout and description are otherwise similar to **Figure 3**.

Nevertheless, SIA showed a slightly smaller overall RMSE (2.75e-3) compared to ImaGene (2.91e-3) (Wilcoxon signed-rank test, *p* = 6.18e-38), and in particular, SIA produces estimates of *s* much closer to 0 for neutral loci. Notably, in this case both SIA and ImaGene were trained with simulations under the same uniform distribution of *s* values (see **Methods**). A different choice of training distribution could impact their performance across selection regimes (see **Discussion**). Furthermore, we binned the results of these methods by both the segregating frequency and the selection coefficient (see **Figure S4**) and again found that in general they exhibit higher variance under low segregating frequency of the beneficial allele. As before, we also tested our regression framework on true and inferred gene trees of test sets simulated under the *S. hypoxantha* demographic model (see **Figure S5**). We found that SIA was approximately unbiased for the moderate (*s* = 0.005) and high (*s* = 0.01) selection regimes but appeared to overestimate the selection coefficient for regions under weak selection (*s* = 0.001 and 0.0025), when both true and inferred genealogies were used as input. Furthermore, SIA appeared to overestimate the selection coefficient for neutral regions when inferred gene trees were used as input, whereas it was approximately unbiased for true gene trees.

### Performance on selection coefficient prediction with different sample sizes

To explore the tradeoffs associated with the use of larger data sets, we examined the performance of SIA under different sample sizes, assuming a constant-sized demographic model (*N*_e_=10,000). **Figure S6** shows the error in selection coefficient inference on a held-out test set, stratified by the age of the allele (panels **A&B**) and present-day derived allele frequency (panels **C&D**) at the site of interest. We observed that sites with low frequency (AF < 0.33) and more recent (onset < 0.2 × 2*N*_e_ generations) alleles experience the most significant reduction in error as sample size increases. Notably, the performance of SIA on more ancient alleles (onset > 0.2 × 2*N*_e_ generations) had little to no improvement as the sample size increased from 32 to 254. These observations are in line with the expectation that having more samples improves the chance of capturing low-frequency alleles, but provides limited information about more ancient events. The reason for this age-dependency is that, looking backwards in time, most lineages coalesce rapidly and only a few survive to more ancient epochs, in a manner that depends only weakly on the sample size. It may be useful to consider these observations when choosing the sample size for use in studying selection in a particular context (see **Discussion**).

### Inference of allele frequency trajectory

We further adapted the deep-learning architecture of SIA to model the allele frequency (AF) trajectory at a site by retaining the output of the LSTM at each time point (**Figure S1**, see **Methods**). We then evaluated the performance of SIA in the inference of the AF trajectory using simulations under the CEU demography across a range of selection coefficients and current DAFs. SIA was largely able to capture the expected trend of more rapidly increasing AF under stronger selection (**Figure S7** and **S9**). In addition, AF estimates by SIA using both true and inferred genealogies were generally unbiased, although AF at more recent time points tended to be slightly underestimated when data was simulated under weaker selection. AF estimates also appeared to be more accurate in terms of variance for alleles under stronger selection (**Figure S8** and **S10**). As expected, the variance of AF estimates tended to increase going further back in time (**Figure S8** and **S10**).

### Model performance on simulations with misspecified demographic models

To evaluate the robustness of SIA to mismatches between the demographic parameters used for simulating training data and the true underlying demography of real data, we tested the method on the selection-coefficient inference task with datasets simulated under a range of alternative parameters. Each aspect of this model misspecification was assessed independently of the others. In particular, the misspecified datasets contained simulations under (*i)* combinations of population mutation (*θ*) and recombination (*ρ*) rates sampled beyond the range used for the training data (**Figures S11** and **S14**), (*ii)* various alternative demographic scenarios (**Figures S12, S15**, and **S17**), and (*iii)* various effective population sizes (**Figures S13** and **S16**). We compared the performance of SIA on these misspecified datasets to that of CLUES [35], supplying both methods with the true genealogies. We consider CLUES the “silver standard” when it comes to robustness because it is unsupervised and therefore should not be susceptible to misspecified training data compared to supervised learning methods such as SIA. Overall, we found that both CLUES and SIA were reasonably robust to model misspecification (**Figures S11-13**), although the performance of both methods inevitably declined when tested on severely misspecified data (**Figure S13**). Interestingly, SIA tended to overestimate selection coefficient when the true *N*_e_ was much smaller than that used for training, and underestimate it when the true *N*_e_ was much larger, whereas CLUES did the opposite (**Figure S13**). Since the CLUES likelihood model of allele frequency transition is parameterized by the population-scaled selection coefficient (*α* = 2*Ns*), a larger *N*_e_ likely appears to CLUES as equivalent to a higher *s*. On the other hand, features used by SIA capture broad information of coalescence and linkage in the ARG, and therefore can be distorted by misspecified *N*_e_ in more subtle ways (see **Discussion**). Using the same misspecified dataset, we also ran SIA with Relate-inferred genealogies and compared its performance to that of the genotyped-based deep-learning model ImaGene [14,15]. In general, SIA appeared to be more robust to model misspecifications, achieving an overall RMSE of 0.00362, 0.00318 and 0.00374 in the misspecified *θ*/*ρ*, demography, and *N*_e_ experiments, respectively, compared to ImaGene, whose RMSE was 0.00416, 0.00330 and 0.00462 in the corresponding experiments (**Figures S14-16**). The advantage of SIA was particularly noticeable in cases of misspecified demographic parameters (**Figures S15 & S16**). Notably, SIA exhibited reduced bias when working with inferred genealogies compared to true genealogies, under conditions of extremely mismatched *N*_e_ (compare **Figures S13 & S16**).

### Model prediction at genomic loci of interest in CEU population

We then applied the SIA model to identify selective sweeps and infer selection coefficients at selected genomic loci in the 1000 Genomes CEU population. These loci included the canonical example of selection at the *MCM6* gene, which regulates the neighboring *LCT* gene and contributes to the lactase persistence trait [36], the *ABCC11* gene regulating earwax production, several pigmentation-related genes, as well as genes associated with obesity, diabetes and addiction (**Table 1**).

**Table 1:**
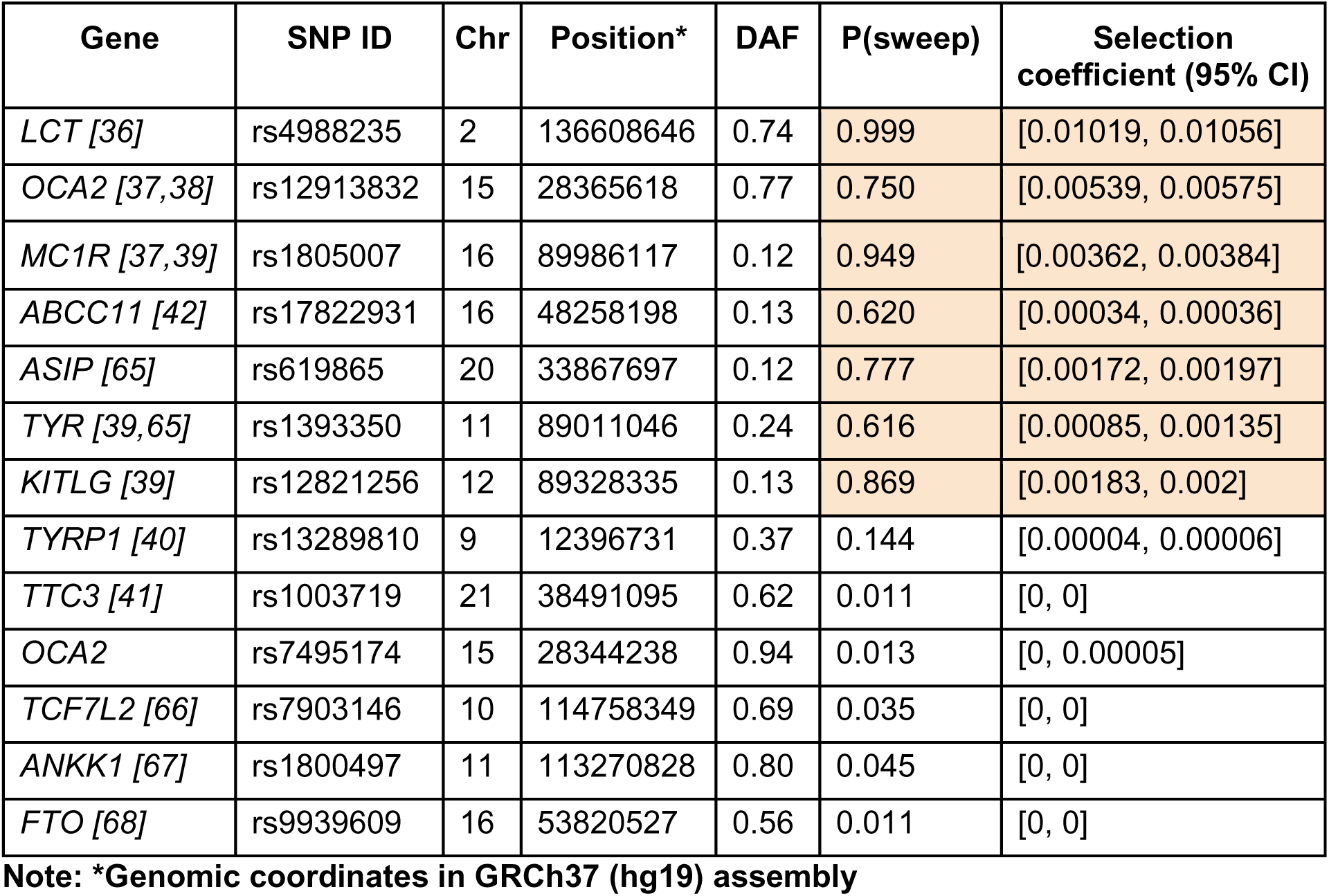
List of genomic loci of interest along with their derived allele frequencies (DAF), sweep probabilities, and selection coefficients inferred by SIA in the 1000 Genomes CEU population.

For *LCT*, SIA detected a strong signal of selection at the nearby SNP that has been associated with the lactase persistence trait (rs4988235). At this SNP, SIA inferred a sweep probability close to 1 and a selection coefficient greater than 0.01, making this one of the strongest signals of selection in the human genome. A close examination of the local genealogy at this site reveals a clear pattern indicative of a selective sweep –– a burst of recent coalescence among the derived lineages (orange taxa are the lineages carrying the derived allele) is clearly visible from the tree (**Figure 5**).

**Figure 5:**
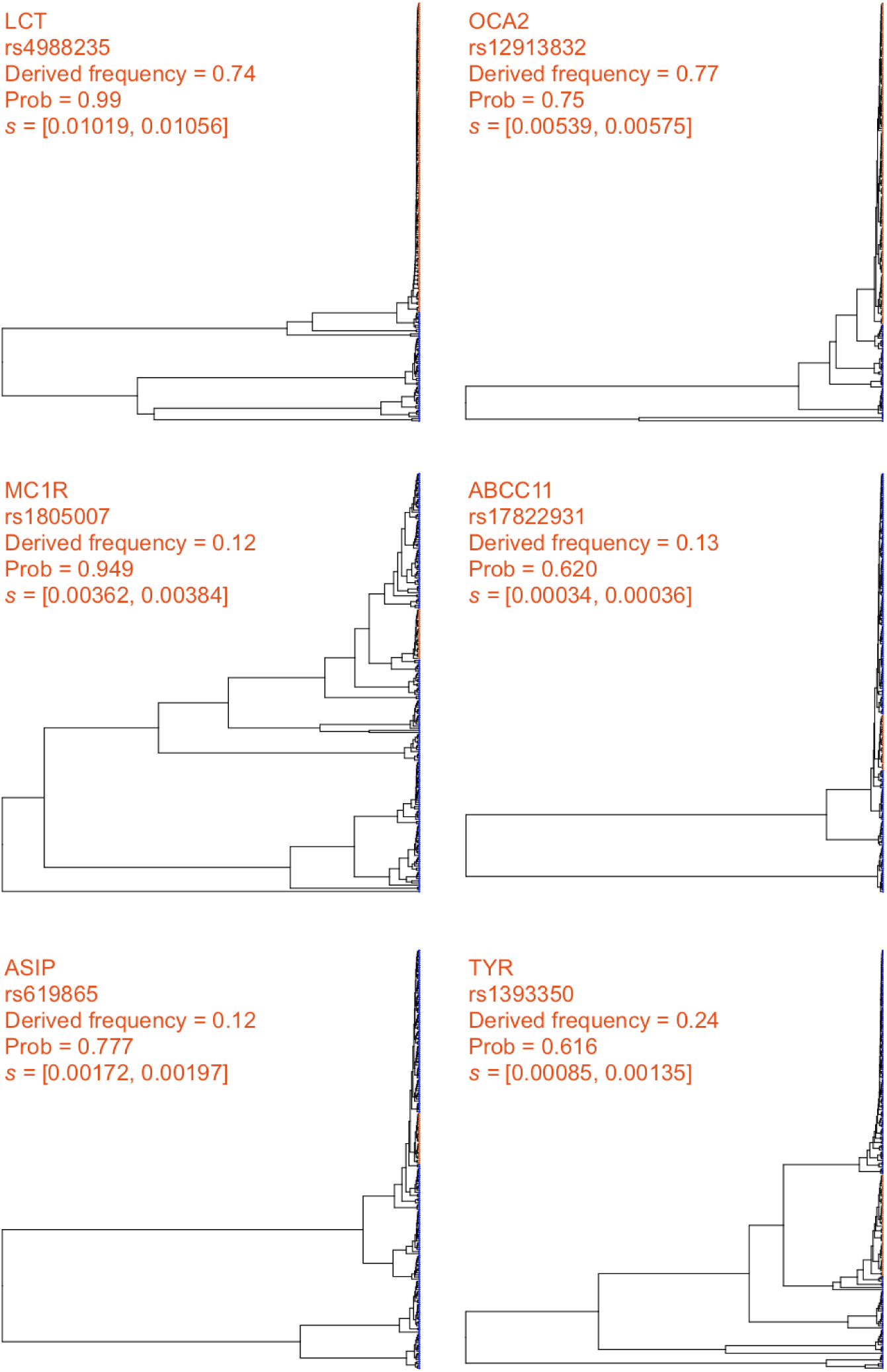
Local genealogies at six loci inferred to be under positive selection in the 1000 Genomes CEU population. Gene name, RefSNP number, derived allele frequency, SIA-inferred sweep probability and SIA-inferred selection coefficient range for each locus are indicated at the top of each panel (see **Table 1** for more details). Taxa carrying the ancestral and derived alleles are colored in blue and orange, respectively.

At a number of pigmentation genes [37–41], SIA detected signals of moderate selection, including *MC1R* (rs1805007, P(sweep) = 0.95, s ≈ 0.0037), *KITLG* (rs12821256, P(sweep) = 0.87, s ≈ 0.0019), *ASIP* (rs619865, P(sweep)= 0.78, s ≈ 0.0019), *OCA2* (rs12913832, P(sweep) = 0.75, s ≈ 0.0056) and *TYR* (rs1393350, P(sweep) = 0.62, s ≈ 0.0011). In addition, SIA identified a weak signal of selection at a SNP in the *ABCC11* gene (rs17822931), which influences earwax and sweat production [42], with a selection coefficient of around 0.00035. There are few other estimates for these genes available for comparison, but, notably, our estimate for *LCT* of *s* ≈ 0.01 is consistent with a previous estimate on the order of 0.01-0.1 [36], and with recent studies of ancient DNA samples [43,44] suggesting a value closer to 0.01. Our estimates suggest that selection at the pigmentation loci is considerably weaker than at *LCT*, in contrast to previous estimates for these loci, which covered a wide range but were generally considerably larger (ranging from 0.02-0.1) [45]. Interestingly, CLUES estimated *s* at the *OCA2* locus to be on the order of 0.001 (roughly similar to SIA’s estimate of 0.0056), but *s* at the *KITLG, ASIP, TYR* loci to be greater than 0.01 (in comparison to SIA’s considerably smaller estimates of 0.0019, 0.0019, and 0.0011) [35]. The apparent discrepancy between the estimates may be partially due to the fact that the two methods used samples from two different populations (CEU for SIA and GBR/British for CLUES).

On the other hand, SIA did not detect significant evidence of positive selection at several disease-associated loci (rs7903146/*TCF7L2*, rs1800497/*ANKK1*, and rs9939609/*FTO*) or at several other pigmentation loci (rs13289810/*TYRP1*, rs1003719/*TTC3*, and rs7495174/*OCA2*) (**Table 1**). Notably, allele frequencies at these six loci tend to be similar in African and European populations [46], suggesting that they are not likely to be under strong environment-dependent positive selection, although it is possible that they have experienced very recent selective pressure that SIA lacks the power to detect (see **Discussion**). Notably, *TYRP1* and *TTC3* also lacked signals of selection in the CLUES analysis. Compared to the genealogies at sweep sites (**Figure 5**), the trees at these putatively neutral loci lack the distinctive signature of recent bursts of coalescence among derived lineages (**Figure 6**).

**Figure 6:**
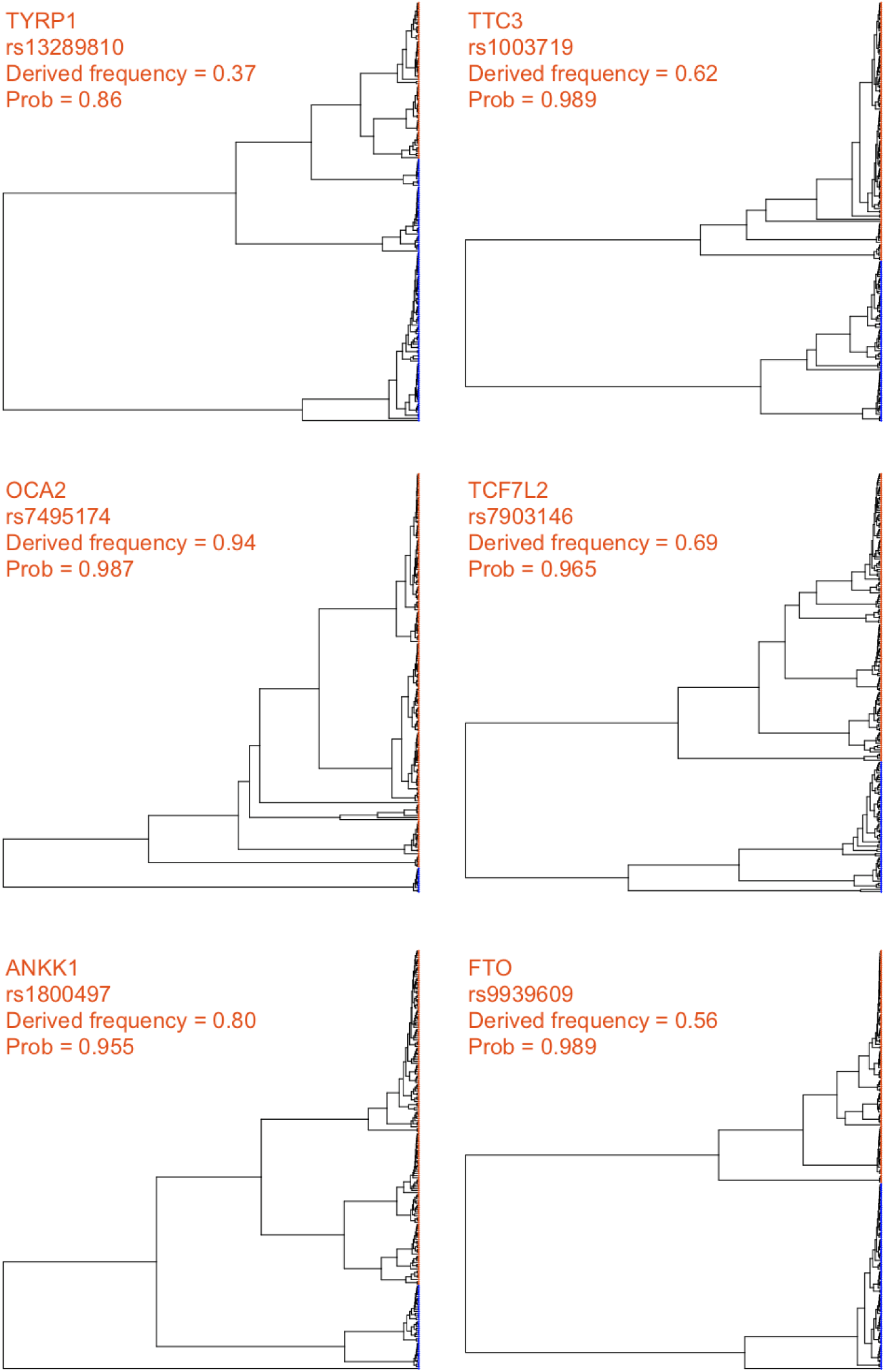
Local genealogies at six loci lacking signal of positive selection in the 1000 Genomes CEU population. Gene name, RefSNP number, derived allele frequency and probability of neutrality inferred by SIA for each locus are indicated at the top of each panel (see **Table 1** for more details). Taxa carrying the ancestral and derived alleles are colored in blue and orange, respectively.

### Southern capuchino species analysis

Our previous study of southern capuchino seedeaters made use of the full ARG and machine learning to detect and characterize selective sweeps, and suggested that soft sweeps are the dominant mode of adaptation in these species (see **Methods** for more details). To further characterize the targets and strengths of positive selection in these species, we applied SIA to polymorphism data [47] for *S. hypoxantha*, and adopted a conservative approach by reporting only sites with DAF ≥ 0.5, SIA-inferred *s* ≥ 0.0025, and SIA-inferred sweep probability ≥ 0.99 (see **Methods**). In addition to loci near top *F*_ST_ peaks and known pigmentation-related genes (**Table 2**), we identified many more sites under positive selection located outside the previously scanned *F*_ST_ peaks, amounting to a total of 15,551 putative partial soft sweep sites across the 333 scanned scaffolds for *S. hypoxantha*. These sites can be prioritized for further evaluation and downstream analysis. Notably, SIA enabled us to distinguish between selection at regulatory and coding sequences, and we found that sweep loci near *F*_ST_ peaks and pigmentation genes fall mostly in non-coding regions (**Table 2**). We additionally surveyed all putative sweep sites identified by SIA and found that they are indeed enriched in non-coding regions (Fisher’s exact test, *p* = 6.80×10^−5^), particularly noticeable in the “near-coding” regions (**Figure S21**). Consistent with the observation that the most highly differentiated SNPs among taxa are non-coding [47,48] our finding suggests that positive selection may act on *cis*-regulatory regions to drive differentiation and the subsequent speciation process. Furthermore, we examined many individual predictions in detail, considering the local trees inferred by Relate at these high-confidence predictions (**Figure 7**). We found, in numerous cases, that these sweeps had distinct genealogical features, displaying evidence of a burst of coalescence events, corresponding to unusually large and young clades. Prominent examples include predictions near pigmentation-related genes *ASIP, KITL, SLC45A2*, and *TYRP1*.

**Table 2:**
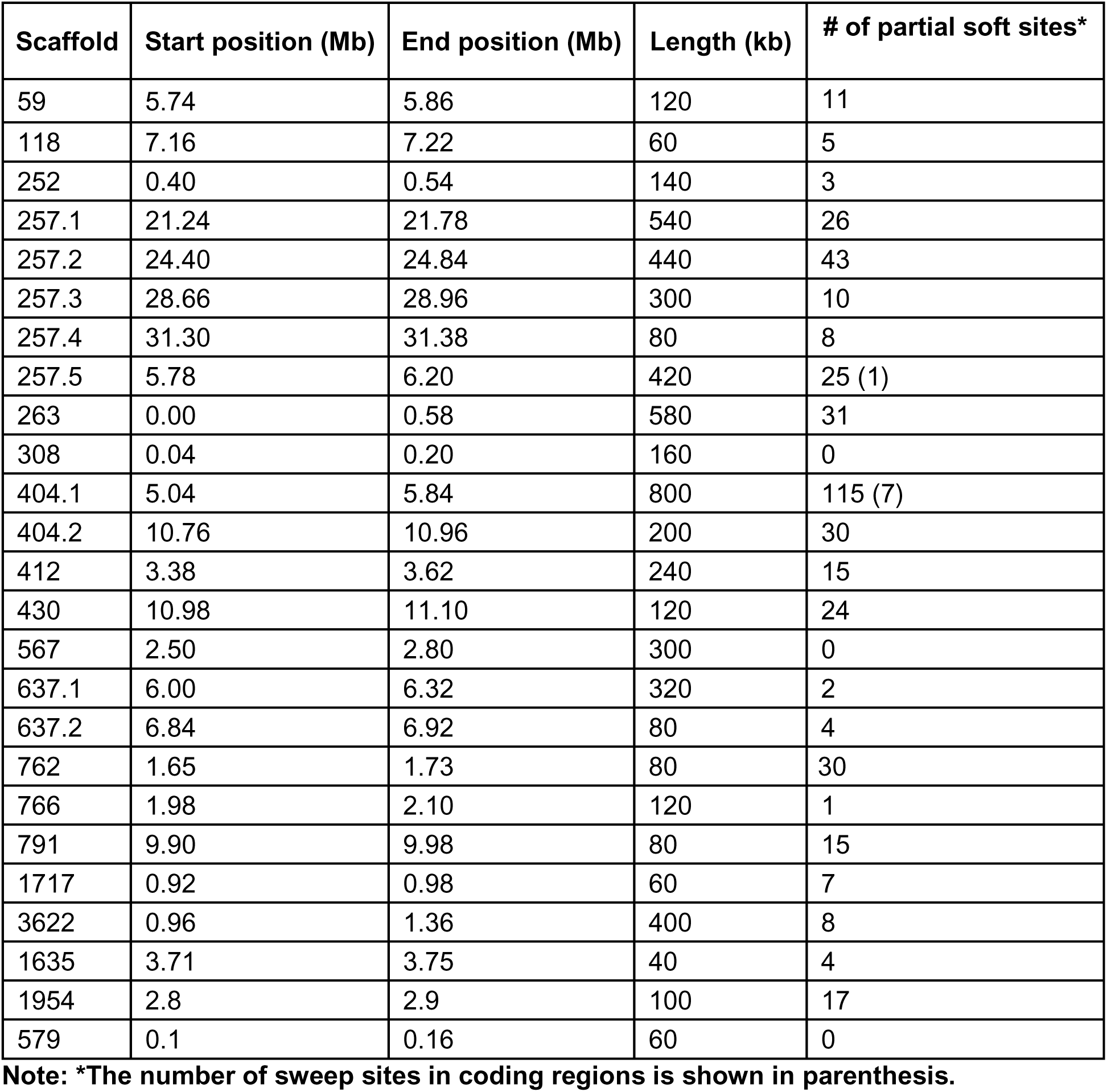
The top 25 *F*_ST_ peaks identified in [31] along with the number of partial soft sites in *S. hypoxantha* identified for each scaffold using SIA. To avoid cases with limited power, we focused on sites with segregating frequency ≥ 0.5, SIA-inferred *s* > 0.0025, and SIA-inferred sweep probability ≥ 0.99.

| Scaffold | Start position (Mb) | End position (Mb) | Length (kb) | # of partial soft sites* |
| --- | --- | --- | --- | --- |
| 59 | 5.74 | 5.86 | 120 | 11 |
| 118 | 7.16 | 7.22 | 60 | 5 |
| 252 | 0.40 | 0.54 | 140 | 3 |
| 257.1 | 21.24 | 21.78 | 540 | 26 |
| 257.2 | 24.40 | 24.84 | 440 | 43 |
| 257.3 | 28.66 | 28.96 | 300 | 10 |
| 257.4 | 31.30 | 31.38 | 80 | 8 |
| 257.5 | 5.78 | 6.20 | 420 | 25 (1) |
| 263 | 0.00 | 0.58 | 580 | 31 |
| 308 | 0.04 | 0.20 | 160 | 0 |
| 404.1 | 5.04 | 5.84 | 800 | 115 (7) |
| 404.2 | 10.76 | 10.96 | 200 | 30 |
| 412 | 3.38 | 3.62 | 240 | 15 |
| 430 | 10.98 | 11.10 | 120 | 24 |
| 567 | 2.50 | 2.80 | 300 | 0 |
| 637.1 | 6.00 | 6.32 | 320 | 2 |
| 637.2 | 6.84 | 6.92 | 80 | 4 |
| 762 | 1.65 | 1.73 | 80 | 30 |
| 766 | 1.98 | 2.10 | 120 | 1 |
| 791 | 9.90 | 9.98 | 80 | 15 |
| 1717 | 0.92 | 0.98 | 60 | 7 |
| 3622 | 0.96 | 1.36 | 400 | 8 |
| 1635 | 3.71 | 3.75 | 40 | 4 |
| 1954 | 2.8 | 2.9 | 100 | 17 |
| 579 | 0.1 | 0.16 | 60 | 0 |

**Note:** \*The number of sweep sites in coding regions is shown in parenthesis.

**Figure 7:**
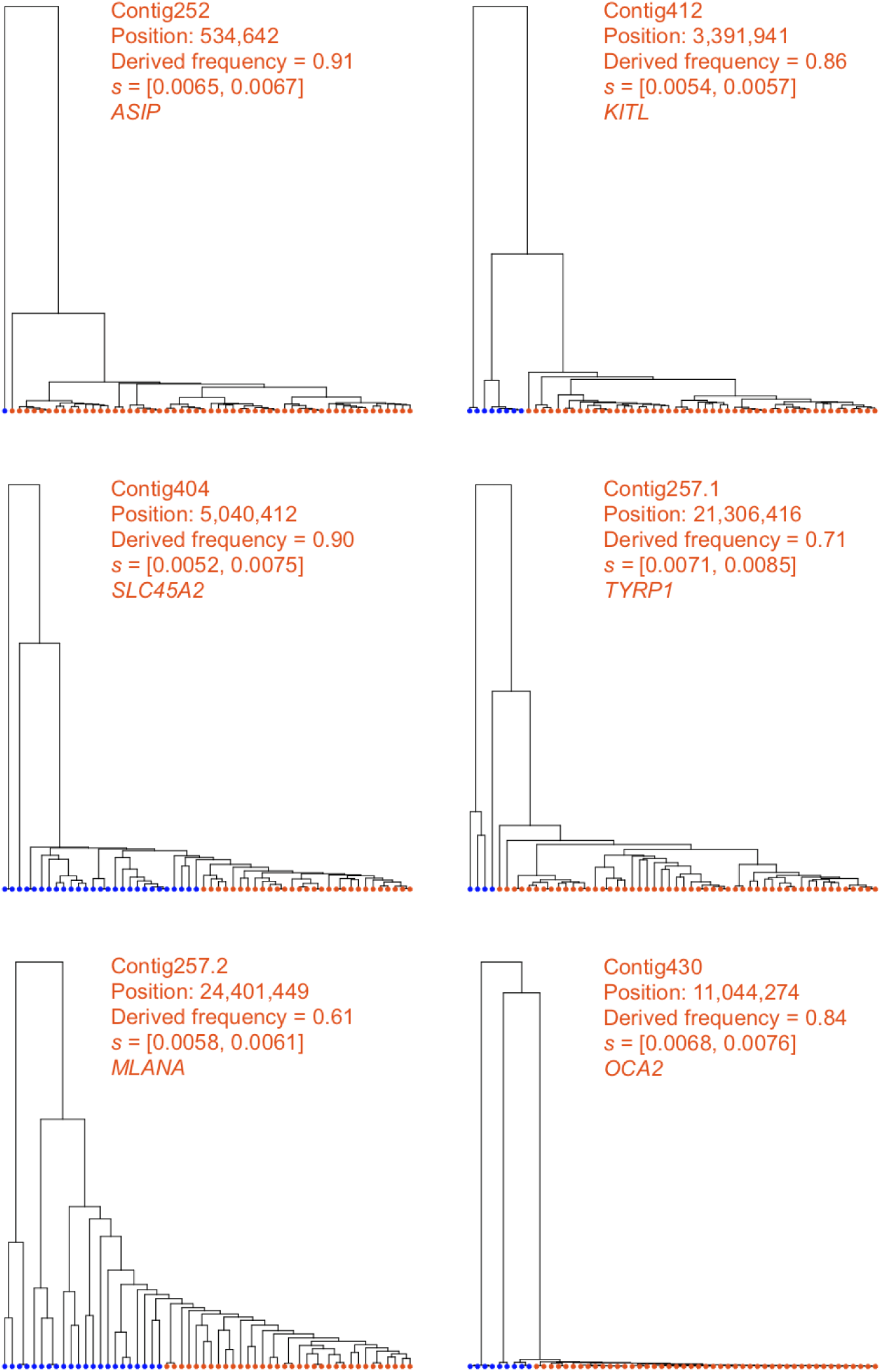
Local genealogies at five loci inferred to be under positive selection in *S. hypoxantha*. Contig name, position of SNP, derived allele frequency, SIA-inferred selection coefficient range, and the pigmentation gene closest to the locus in question are indicated at the top of each panel. Haploid genomes carrying the ancestral and derived alleles are colored in blue and orange, respectively.

## Discussion

The ARG is useful for addressing a wide variety of biological questions ranging from inferring demographic parameters to estimating allele ages. SIA exploits the particular utility of the ARG for accurate inference of positive selection in a way that makes use of the full dataset, as opposed to traditional summary statistics, which necessarily discard substantial information. Direct use of the ARG improves upon traditional summary statistics in two key ways. First, it enables consideration of the temporal distribution of coalescence and recombination events in the history of the analyzed sequences, in contrast to traditional summary statistics that simply average over these coalescence and/or recombination events. In addition, ARG-based methods provide better spatial resolution by separately examining individual genealogies and the recombination breakpoints between them, rather than averaging across windows containing unknown numbers of genealogies. These detailed patterns of coalescences and linkage enable the ARG-based approaches to capture a more localized and fine-grained picture of selection (e.g. infer selection coefficient and allele frequency trajectory) as well as to achieve a better classification performance. This performance advantage is particularly noticeable at lower DAFs and when selection is weak, a regime where previous methods for selection inference fall short (**Figure 2**).

At the same time, the supervised machine-learning approach sets SIA apart from another ARG-based method, CLUES, which approximates a full likelihood function for ARGs in the presence of selection using importance sampling and a HMM. Although the accuracy of both SIA and CLUES degraded when using inferred genealogies compared to true genealogies, reflecting the error and uncertainty at the ARG inference step, SIA appeared to be more robust to gene tree uncertainty (**Figures 3** and **4**). One possible reason for this observation is that CLUES effectively assumes that the selection coefficient at the focal site is conditionally independent of the flanking trees given the focal tree. This assumption should hold in the presence of fully specified genealogies, but it may make CLUES more sensitive to errors in the inferred genealogies. In other words, through its use of supervised learning, SIA may be able to compensate for the effects of genealogy inference error on its estimation of the selection coefficient by also directly considering the flanking trees and LD-related patterns among them. Still, the drop in accuracy observed across methods underscores the dependency of ARG-based approaches on the ARG inference method. For this reason, we anticipate that SIA may benefit substantially from further improvement in ARG inference tools (see ref. [9]).

The ARG-based feature set distinguishes SIA from other supervised machine learning approaches for characterizing selective sweeps. SIA uses local topological features of the ARG, which are more informative than the SFS-or LD-based summary statistics employed by machine learning methods such as S/HIC, SFselect, and evolBoosting. Using simulations, we demonstrated that the SIA classifier outperformed a deep-learning method that aggregates these traditional summary statistics (**Figure 2**). We also compared SIA with ImaGene, which represents another flavor of supervised learning methods, inspired by the recent rise of CNNs for image recognition. ImaGene encodes sequence alignments as images for powerful population genetic inferences with CNNs and provides a state-of-the-art benchmark to compare against. We found that ImaGene performs remarkably well across a wide range of simulations, but SIA does appear to be somewhat less biased and more robust to model misspecification than ImaGene. The evolutionary information in the ARG is implicit in the sequence alignment but some of this information may be difficult for a brute-force machine learning model to discover directly.

We demonstrated that utilizing the ARG granted SIA considerably improved performance over deep learning models solely employing traditional summary statistics. However, a possible drawback of an ARG-based model is the potentially prohibitive computational overhead incurred by ARG inference, especially as sample size grows. Picking a sample size when running SIA involves a tradeoff between scalability (fewer samples, faster ARG inference) and performance (more samples, slower ARG inference). We have found that SIA can infer selection coefficients reasonably well with as few as 16 haplotypes. Including more samples did improve performance but with a sublinear reduction in error (**Figure S6**). Therefore, a sample size from a few dozen to a few hundreds — well within the capabilities of most modern ARG inference methods — strikes a good balance between performance and scalability. Moreover, we found that larger sample sizes improved prediction performance primarily for alleles at lower frequencies but had little impact on the performance for more ancient alleles (as most lineages would have already coalesced going further back in time) (**Figure S6**). This observation suggests that the choice of the sample size when applying SIA should be guided by the biological question of interest –– ancient selection can be studied with just a handful of samples, whereas a larger sample size is better suited to detect more recent sweeps.

Like other supervised learning methods, SIA relies on simulations to generate training data, and therefore could be biased by subjective choices of simulation parameters. For example, SIA and ImaGene cannot make accurate predictions of selection coefficients outside the range represented in the training data (**Figure S18**), whereas unsupervised methods such as CLUES are not limited to a pre-defined range (**Figure S19**). This problem could be circumvented by training on an extended range of *s*. Similarly, the tendency of SIA to underestimate the selection coefficient for sites under weak selection (**Figures 3, 4**) could be mitigated by augmenting the training set with simulations densely sampled from the weak selection regime. A more subtle issue, however, arises when the underlying generative process of the real data does not match the assumptions made for the simulations of the training data, potentially compromising the accuracy of the method when applied to real data. Thus, we tested SIA on simulations with parameters mismatching those used in the training procedure. In general, we found that SIA was fairly robust to alternative parameter values, although, as expected, performance did degrade somewhat under severely misspecified models. Notably, SIA achieved a similar level of robustness to model parameter misspecification as the unsupervised (i.e. not relying on training data) likelihood method CLUES, yet outperformed the supervised deep learning method ImaGene.

Applying SIA to the CEU panel from the 1000 Genomes Project yielded several noteworthy findings at loci with known ties to phenotypes of interest. In addition to confirming the canonical signal of selective sweep at the *LCT* locus, SIA detected a novel signal of selection at a GWAS SNP in the *MC1R* gene associated with red hair color, contrasting a previous study that could not find evidence of selection at *MC1R* in the European population [49]. The derived allele at this locus segregates at around 10% in the CEU population but is nearly absent in non-European populations [46]. In addition, at the *MC1R* locus the Relate test statistic for selection [25], which tends to perform particularly well at low segregating frequencies (**Figure 2**), falls slightly below the significance threshold of 0.05, supporting the evidence of positive selection at this locus. SIA also detected evidence of selection at a SNP in the *ABCC11* gene reported to be the determinant of wet versus dry earwax as well as sweat production, mirroring the signal of selection previously found in the East Asian population [50], although selection in the CEU population appeared to be much weaker. In addition, SIA identified selection at a few other pigmentation-related loci, yet determined previously identified SNPs in the *TYRP1* and *TTC3* genes to be largely free from selection (**Table 1**). These results were consistent with a previous study [35], which reported similar results for these pigmentation-related loci, albeit in a slightly different population (GBR). SIA notably did not detect positive selection at GWAS loci in the *TCF7L2* gene associated with type-2 diabetes, the *ANKK1* gene implicated in addictive behaviors, and the *FTO* gene associated with obesity. Overall, this empirical study with the 1000 Genomes CEU population has illustrated how SIA can be applied to assess natural selection at the resolution of individual sites, suggesting that it may be useful in prioritizing GWAS variants for further scrutiny.

In our previous work on southern capuchino seedeaters [31] (see **Methods**), we applied newly developed statistical methods for ancestral recombination graph inference and machine-learning for the prediction of selective sweeps. We found evidence suggesting that a substantial fraction of soft sweeps are partial but had limited power to identify them (i.e. average accuracy of 56%). SIA considerably improved our characterization of positive selection in the southern capuchino species in two key ways. The SIA framework performs inference of selection directly from genealogies instead of traditional summary statistics, and in doing so achieved an accuracy of up to 96% in detecting partial soft sweeps. Consequently, we found abundant evidence of soft sweeps beyond the previously scanned *F*_ST_ peaks, and additionally were able to estimate their selection coefficients. Importantly, SIA also took the analysis of selection beyond broad genomic windows containing sweeps to the identification of specific putative causal variants. We took advantage of this substantial improvement in genomic resolution and analyzed the distribution of these sweep sites, which revealed that positive selection on regions that likely contain *cis*-regulatory elements plays a role in driving the differentiation and speciation of southern capuchino seedeaters.

While we believe SIA represents an important step forward in the use of the ARG for machine-learning-based selection inference, there remain several possible avenues for improvement. For example, SIA currently uses a point-estimate of the ARG, rather than a distribution, and therefore does not explicitly take gene-tree uncertainty into account. We plan to improve SIA by using strategies for inferring approximate posterior distribution of ARGs (e.g., [24]), as well as designing better algorithms for ARG reconstruction that balance accuracy with scalability and can handle thousands of genomes. In addition, the SIA framework was applied in the context of single-locus selective sweeps, but could be extended to study polygenic selection, by making use of summary statistics from genome-wide association studies (as in [51]) and adapting the architecture of our neural network to account for selection acting at multiple sites. Finally, the robustness of SIA to model misspecifications can be further improved by ensuring the simulated data is generated under a distribution that is compatible with the real target data set. We anticipate that the continual advancement in ARG inference methods has the potential to open up many new applications for this flexible and powerful model of ARG-based deep learning in population genetics.

## Methods

### Simulated datasets used for training and testing the selective sweep model

Training and testing data sets were generated using discoal [52] by simulating 1,000,000 regions of length 100 kb for each model we considered (i.e., “neutral” or “hard sweep”). Aside from these regions, 2,000 were simulated for validation and 5,000 were simulated for testing. The number of sampled sequences was selected to match the number of individuals in the CEU population in the 1000 Genomes dataset. Thus, a total of 198 haploid sequences were sampled. Simulations used a demographic model based on European demography [34]. In non-neutral simulations, selection was applied to a single focal site located in the middle of the simulated region. We sampled each of the main demographic and selection parameters from a uniform distribution: (1) mutation rate *μ* ∼ *U*(1.25e-08, 2.5e-08), (2) recombination rate *ρ* ∼ *U*(1.25e-08, 2.5e-08), (3) selection coefficient *s* ∼ *U*(0.0001, 0.02), and (4) segregating frequency of the site under selection *f* ∼ *U*(0.01, 0.99).

### ARG Feature Extraction

For each target variant, we extracted the corresponding gene tree from the ARG, then overlaid it with 100 discrete timepoints. These timepoints were fixed across all trees in an approximately log-uniform manner that resulted in finer discretization of more recent time scales (as in [24]). We considered biallelic sites only and assumed no recurrent mutations; thus each mutation was assumed to occur on the branch of the tree where the ancestral allele switches to the derived. For each timepoint, we calculated the number of active ancestral and derived lineages. Furthermore, we computed the number of all active lineages (not distinguishing between ancestral and derived) at the same set of predefined timepoints in the two left and right flanking gene trees to account for linkage disequilibrium. Together, these features were summarized in a 600-dimensional feature vector, which was then used as input to an RNN. The feature of a simulated sweep region was extracted from the sweep site (by default at the center in all simulations) whereas the feature of a simulated neutral region was extracted from a variant site (randomly chosen) with a pre-defined matched derived allele frequency. The features for each genomic locus of interest in the CEU population were extracted from all variant sites at that locus having a derived allele frequency of >0.05.

### Training an RNN to predict different modes of selection

An RNN was applied to the simulated training data sets to learn a classification or regression model for the task at hand. We used a Long Short-Term Memory (LSTM), a particular form of RNN, to accommodate the temporal nature of our features and account for long-term dependencies and the vanishing gradient problem observed in traditional RNNs. Our model had 100 timepoints with the final target output depending on the use of classification or regression. For the classification task, the final target output is a label for a binary classification problem predicting whether a region is under selection or neutrality. For the regression task, the final target output is a continuous value, representing the selection coefficient or the time of selection onset. We also took a many-to-many approach to model the allele-frequency trajectory for the site under selection. The *Keras* software was used to train and test the model. We used a two-stacked LSTM to account for greater model complexity where the number of units in each stack was set to 100 and the hyperbolic tangent (*tanh*) was used as an activation function. The *Adam* optimization method with its default operating parameters was used to update the network weights. For the classification task, the *Softmax* activation function was applied on the final dense layer and the *binary_crossentropy* was used to compute the cross-entropy loss between true labels and predicted labels. For the regression task, the *linear* activation function was applied on the final dense layer and the *mean_squared_error* function was used.

### Estimation of Confidence Intervals

To turn our single-valued regression model into one capable of returning a distribution of predictions of *s*, we reused the dropout technique that is typically used during training. Dropout enables a fraction of nodes to be randomly “turned off” in a certain layer, which assists in the regularization of the model and helps prevent overfitting. We applied dropout during inference, enabling us to sample a “thinned” network to generate a sample prediction. By repeatedly sampling thinned networks, we generated a distribution of predictions and then computed confidence intervals based on this distribution [53].

### ARG Inference

Relate [25] (v1.0.17) was used for inferring ARGs underlying simulated genomic samples as well as the CEU population in the 1000 Genomes dataset. For simulations under the Tennessen *et al*. demography [34], Relate was run with the true simulation parameters (*μ, ρ* and *N*_*e*_) specified; whereas for genomic loci of the CEU population, Relate was run with a mutation rate of 2.5×10^−8^ /base/generation (-m 2.5e-8), a constant recombination map of 1.25×10^−8^/base/generation and a diploid effective population size of 188,088 (-N 376176). The choice of mutation rate follows [35] based on estimates from [54]. Although some more recent estimates have been lower [55], these differences in mutation rate are unlikely to have a major effect on our selection inference since SIA appears to be fairly robust to misspecification of mutation rate (**Figures S11 & S14**). For simulations and genomic loci of the *S. hypoxantha* population, Relate was run with *μ*=*ρ*=1×10^−9^ /base/generation and a diploid *N*_*e*_ of 130,000. The branch lengths of Relate-inferred genealogies were estimated iteratively with the ‘*EstimatePopulationSize*.*sh*‘ script in the Relate package. Specifically, population size history was inferred from the ARG, the branch lengths are then updated for the estimated population size history and these steps are repeated until convergence. This was done for a default of 5 iterations (--num_iter 5).

### Alternative methods for selection inference

To benchmark the performance of SIA for classification of sites under neutrality versus selective sweep, we ran the following methods: Tajima’s D [10], H1 [32], iHS [33], a summary statistics-based deep learning model, and a tree-based statistic that is part of the Relate [25] program. Tajima’s D, H1 and iHS were calculated with the *scikit-allel* package. Haplotypes of the entire 100kb simulated genomic segment were used for Tajima’s D and H1 calculations. The unstandardized iHS was computed at every site with minor allele frequency > 5%, with respect to all other sites in the genomic segment (min_maf=0.05, include_edges=True). iHS scores of all sites were then standardized in 50 allele-frequency bins. Finally, the iHS score of a genomic region was taken to be the mean of the iHS scores of all of its variant sites. For the summary statistics-based deep learning model, we made use of the summary statistics used by S/HIC [12,13] as features for our deep learning architecture. These included 11 sequence-based summary statistics (see **Figure 3** in [56]) which were used as features for our deep learning model to distinguish among the two classes at hand (selective sweep versus neutral drift). All statistics were collected along five consecutive 20-kb windows with the objective of identifying possible sweeps induced by a positively selected mutation in the third (middle) window. Some of these summary statistics corresponded to standard measures of diversity, such as ss (the number of segregating sites), π [57], Tajima’s D [10], θ_W_ [58], θ_H_ [59], the number of distinct haplotypes [60], H1, H12, H2/H1 [32], Z_nS_ [61], and maximum value of ω [62]. For each of these statistics, we computed an average value for each of the five 20 kb windows for the simulated population. Finally, each summary statistic was normalized by dividing the value recorded for a given window by the sum of values across all five windows. The Relate tree-based selection test was performed with an add-on module (*DetectSelection*.*sh*) using the inferred genealogy with calibrated branch lengths at a site of interest, yielding a log_10_ p-value for each site. We also compared the performance of SIA for selection coefficient inference to that of CLUES [35] and a genotype-based convolutional neural network (CNN) framework [14,15]. Selection coefficient inference from true genealogies was performed with *clues-v0* (https://github.com/35ajstern/clues-v0). Transition probability matrices were built on a range of selection coefficients [0, 0.05] at increments of 0.0001 and present-day allele frequencies [0.01, 0.99] at increments of 0.01. Selection-coefficient inference from Relate inferred genealogies was performed with CLUES (https://github.com/35ajstern/clues). Branch lengths of the genealogy at the site of interest were resampled with Relate for 600 MCMC iterations, and CLUES was run with the following arguments: ‘--tCutoff 10000 --burnin 100 --thin 5‘. For the genotype-based CNN model, each simulated genomic segment was preprocessed by first sorting the haplotypes and then converting the segment to a fixed-size genotype matrix. Haplotype sorting was performed by *1)* calculating the pairwise manhattan distances between haplotypes, *2)* setting the haplotype with the smallest total distance to all other haplotypes as the first haplotype, and *3)* sorting the remaining haplotypes in increasing distance to the first haplotype. To convert the sorted haplotypes to a fixed-size genotype matrix, centered on the middle variant of a simulated region, up to 180 variants on each side were retained. Variants beyond 180 were discarded and if there were fewer than 180, the missing variants were padded with zeros. Ancestral and derived alleles were coded with 0’s and 1’s, respectively. Consequently, each simulated genomic region was encoded as a (198 × 360) binary matrix, along with a real-valued vector encoding the genomic positions of the variants in the matrix. The CNN model had a branched architecture –– one branch with five 1D convolution layers taking the genotype matrix as input and another branch with a fully connected layer taking the vector of variant positions as input. The output of the two branches was flattened, concatenated and fed into 3 fully connected layers, followed by a linear output layer to predict selection coefficient (**Figure S20**).

### Evaluation metrics

To evaluate the performance of SIA’s classification model and alternative methods, we computed a receiver operating characteristic (ROC) curve for the binary class at hand (“neutral” or “sweep”), to provide a more complete summary of the behavior of different types of errors. We further assessed the performance of SIA and alternative methods in terms of correctly predicting the selection coefficient numerically using mean absolute error (mae), root mean square error (rmse), coefficient of determination (*r*^2^), and visually using a box plot that compares the simulated ground truth against the predictions by the method at hand.

### Robustness study

We carried out an extensive analysis of the robustness of our approach, considering not only alternative demographic parameters (such as population size), but also alternative parameters for recombination rate, mutation rate, time of selection onset, and selection coefficients. In all cases, we took care to test our prediction methods under parameters well outside the range used in training.

### Analysis of CEU population in 1000 Genomes data

We applied SIA to infer selection coefficients and allele frequency trajectories in the 1000 Genomes [63] CEU population at 13 genomic loci with known association to phenotypes, some of which were previously identified as likely targets of positive selection (**Table 1**). For each gene of interest, the ARG was inferred with Relate from SNPs within a 2Mb window centered at the gene. Once the ARG was inferred, only SNPs with valid ancestral allele (‘AA‘ INFO field in the vcf file) were retained for downstream analysis. Following the aforementioned protocol (see *ARG feature extraction*), features at all variant sites in the 2Mb window above a derived allele frequency threshold of 0.05 were extracted. Lastly, the SIA model was applied to classify neutrality versus selection, and infer selection coefficient and allele frequency trajectory at each site.

### Localizing sweeps in southern capuchino seedeaters

We recently applied a combination of ARG inference and machine-learning methods for identifying selective sweeps to study previously identified “islands of differentiation” in southern capuchino seedeaters and distinguish among possible evolutionary scenarios leading to their formation [31]. Taking advantage of its improved power and genomic resolution, we applied SIA to sequence data for the species for which we have the most samples, *Sporophila hypoxantha*. We simulated training (250,000 neutral; 250,000 soft sweeps), validation (1000 neutral; 1000 soft sweeps), and testing (2,500 neutral; 2,500 soft sweeps) data sets for SIA under a demographic model inferred by G-PhoCS [64]. Simulations were performed using discoal with the following parameters: (1) mutation rate *μ* = 1e-9, (2) recombination rate *ρ* = 1e-9, (3) derived *N*_e_ = 130,000, (4) root divergence time = 1,850,000 generations ago, (5) root *N*_e_ = 1,450,000, (6) ancestral divergence time = 44,000 generations ago, (7) ancestral *N*_e_ = 14,380,000, (8) selection coefficient *s* ∼ *U*(0.001, 0.02), (9) initial frequency at which selection starts acting on the allele *f*_init_ ∼ *U*(0.01, 0.05), and (10) segregating frequency of the site under selection *f* ∼ *U*(0.25, 0.99). A total of 56 haploid sequences were sampled from each simulation, matching the number of *S. hypoxantha* individuals (28) in the real data. The SIA model for *S. hypoxantha* was built, trained and evaluated in an otherwise similar fashion to that for the CEU population as outlined above.

Using a subset of polymorphism data from [47] of 28 *S. hypoxantha* and 2 *S. minuta* individuals, we applied our trained model to localize selective sweeps in *S. hypoxantha* on 19 scaffolds that contain top *F*_ST_ peaks in at least one pairwise species comparison [48] and/or harbor known pigmentation-related genes such as *ASIP* (located on scaffold 252; induces melanocytes to synthesize pheomelanin instead of eumelanin), *KITL* (located on scaffold 412; stimulates melanocyte proliferation), *SLC45A2* (located on scaffold 404; transports substances needed for melanin synthesis), and CAMK2D (located on scaffold 1717; cell communication), as well as 316 scaffolds that i) are longer than 100kb, ii) contain more than 1,000 variants, and iii) where more than 95% of sites have a consensus ancestral allele, as determined by four identical haplotypes for two individuals from the outgroup species *S. minuta*. The ARG was inferred with Relate for each scaffold independently. Once the ARG was inferred, the SIA model was applied to sites with consensus ancestral allele for classification and selection coefficient inference.

## Acknowledgments

The authors would like to acknowledge Noah Dukler for help with Figure 1 preparation. This research was supported by US National Science Foundation grant (NSFDEB) 1555769, US National Institutes of Health grant R35-GM127070, the CSHL School of Biological Sciences Gladys & Roland Harriman Fellowship, and the Simons Center for Quantitative Biology at Cold Spring Harbor Laboratory. The content is solely the responsibility of the authors and does not necessarily represent the official views of the US National Institutes of Health or the US National Science Foundation.

## Availability of data and materials

The scripts used for analyses in this study are available at github.com/CshlSiepelLab/arg-selection under a GNU GPLv3 license.

## Notes

### Competing Interest Statement

The authors have declared no competing interest.

