## Supplemental figures for "SIA: Selection Inference Using the Ancestral Recombination Graph"

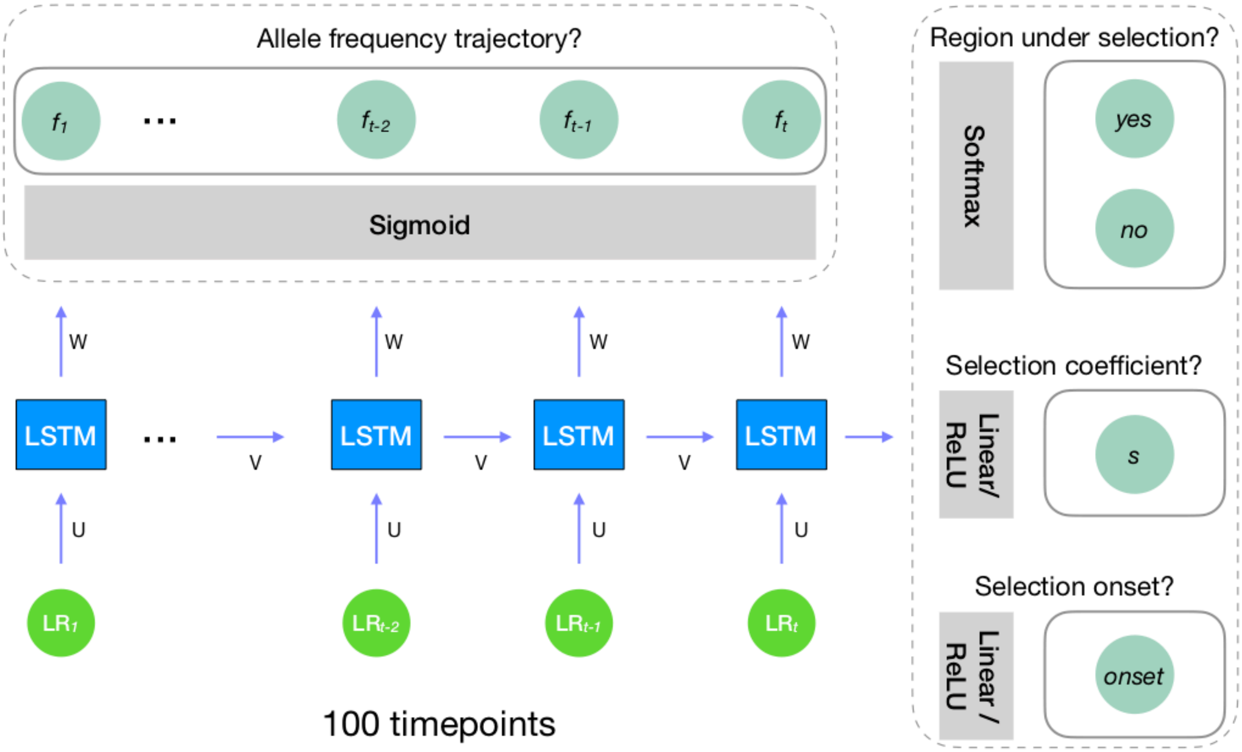


**Figure S1: Overview of the deep learning architecture.** A form of Recurrent Neural Networks called Long-Short Term Memory (LSTM) was used for sweep prediction. LSTMs are designed to handle the temporal nature of our feature set and account for long term dependencies. Our model has 100 timepoints with the final target output differing in terms of the task at hand (i.e. classification or regression task). For the classification task, the final target output is a label for a binary classification problem predicting whether a region is under selection or neutrality. For the regression task, the final target output is a continuous value, representing the selection coefficient or the selection onset. We also took a many-to-many approach to model the allele frequency trajectory for the site under selection.

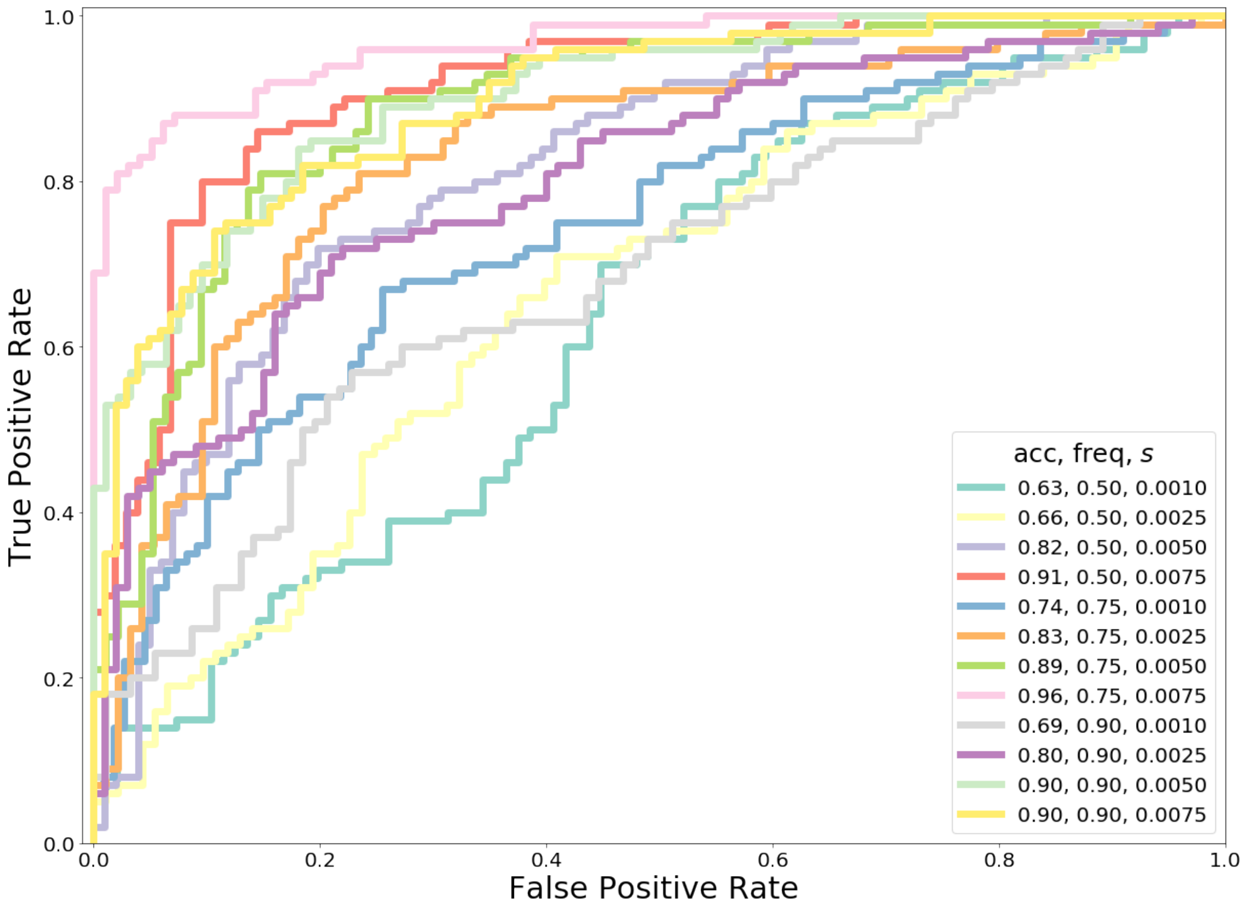


**Figure S2: The impact of selection on the performance of SIA on the *S. hypoxantha* population.** Data was simulated under a variety of selection regimes and segregating frequencies for the beneficial mutation under selection (shown in the legend under freq and s). The prediction task involves two classes: neutral versus soft sweep. SIA was tested on a set of 200 regions per ROC curve (100 per class), and the receiver operating characteristic (ROC) curve records the true positive rate (TPR) as a function of the false positive rate (FPR). The curve associated with the binary prediction task (neutral vs. soft sweep) is obtained by varying the prediction threshold from 0 to 1 and recording for each threshold the number of regions correctly assigned (TPs) and misassigned (FPs) (with prediction probability above the threshold). The performance of SIA was evaluated based on the area under its ROC curve, or AUROC (shown in the legend under acc). We report SIA's AUROC as an average across 200 replicate datasets for each ROC curve.

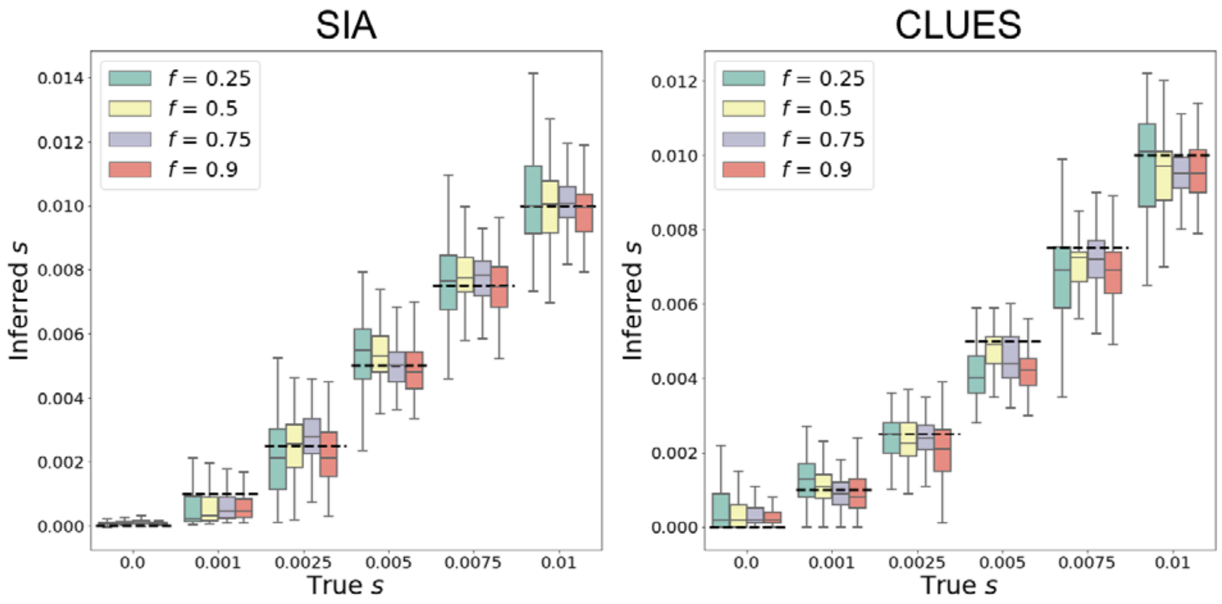


**Figure S3: Predictions of selection coefficient on simulations using true gene trees.** Results are binned by segregating frequency for each selection regime. Each model condition (i.e. box plot) represents a set of 100 replicates. Figure layout and description are otherwise similar to **Figure 3**.

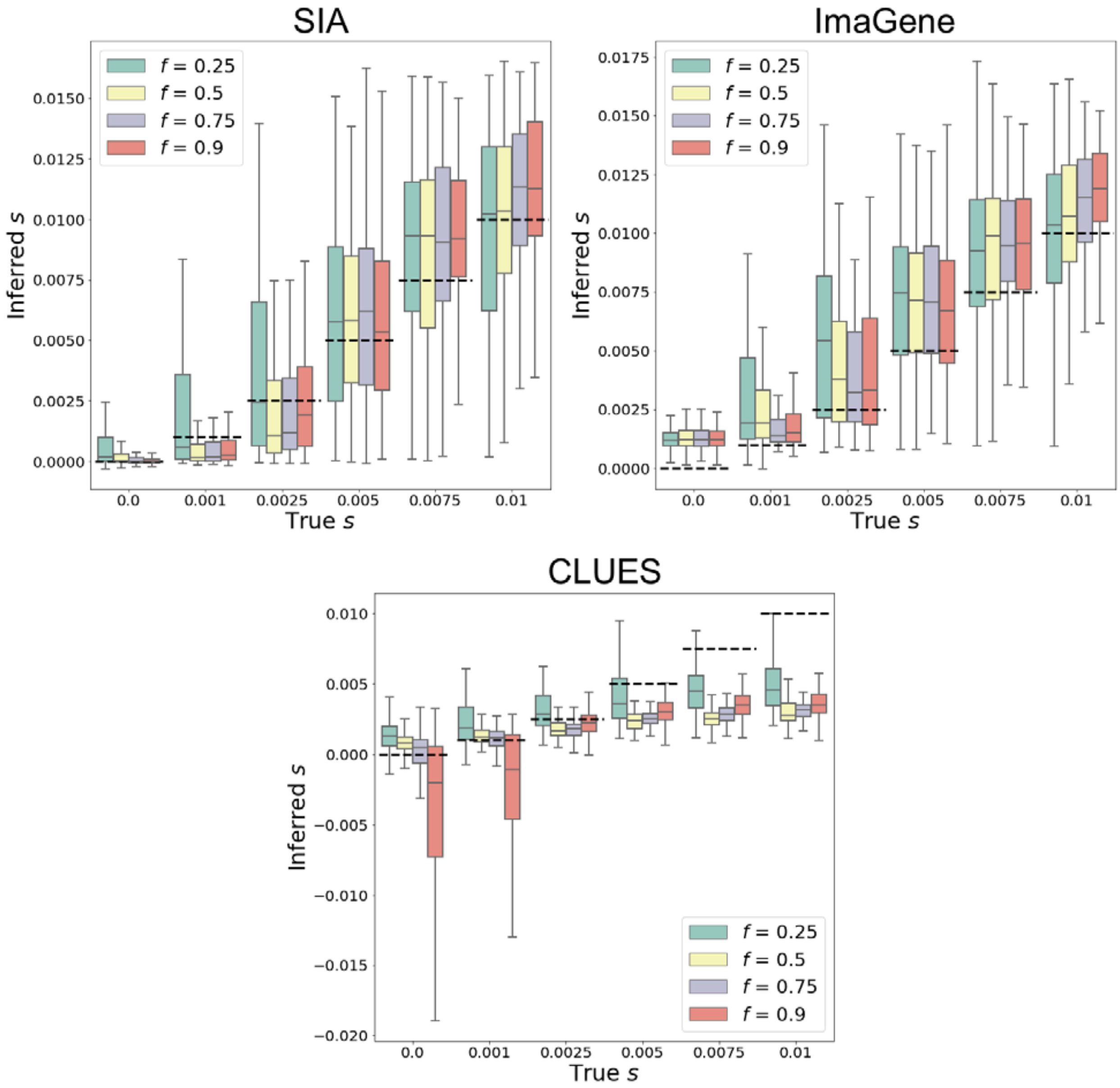


**Figure S4: Predictions of selection coefficient on simulations using SIA, ImaGene, and CLUES.** Results are binned by segregating frequency for each selection regime. Each model condition (i.e. box plot) represents a set of 100 replicates. The simulations are based on the CEU demographic model where inferred genealogies were used as input to SIA and CLUES. Figure layout and description are otherwise similar to **Figure 3**.

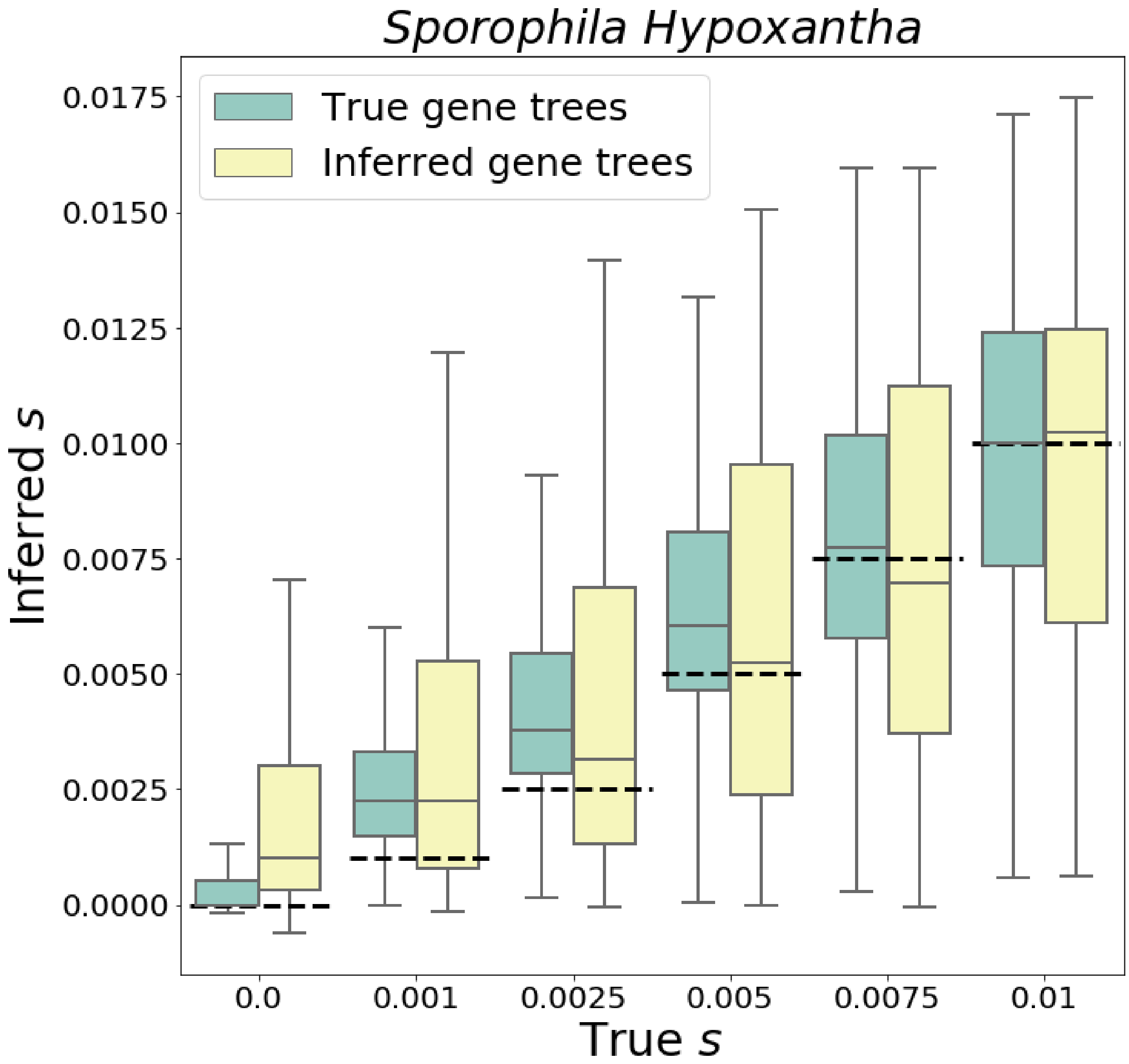


**Figure S5: Predictions of selection coefficient on southern capuchino simulations using SIA.** The distribution of inferred selection coefficients for SIA on *S. hypoxantha* and each model condition is reported using a box plot. The simulations are based on the capuchinos demographic model where true or inferred genealogies were used as input to SIA. Each model condition (i.e. box plot) represents a set of 400 replicates. Figure layout and description are otherwise similar to **Figure 3**.

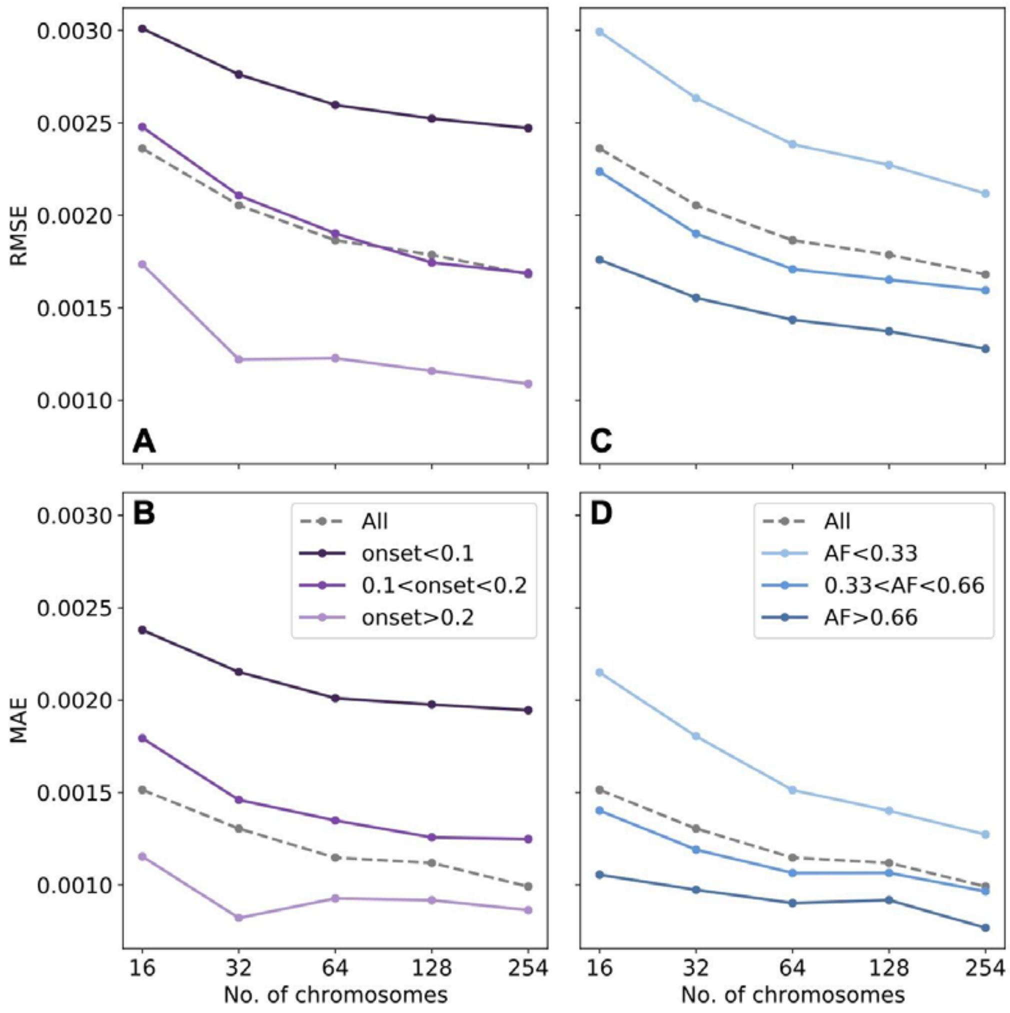


**Figure S6: Performance of SIA in selection-coefficient inference with different sample sizes.** A separate SIA model was trained using 100,000 neutral and 100,000 sweep simulations (95%-5% train-validation split) under a constant $N_e$ = 10,000 for 16, 32, 64, 128 and 254 haploid genomes. The performance of each model in selection-coefficient inference was evaluated on a test set of 10,000 neutral and 10,000 sweep simulations using root mean square error (RMSE, top) and mean absolute error (MAE, bottom), stratified by time of emergence (in coalescent unit of $4N_e$) of the *de novo* beneficial allele (left), and by the current derived allele frequency (right). Grey dots indicate the overall performance on the entire test set.

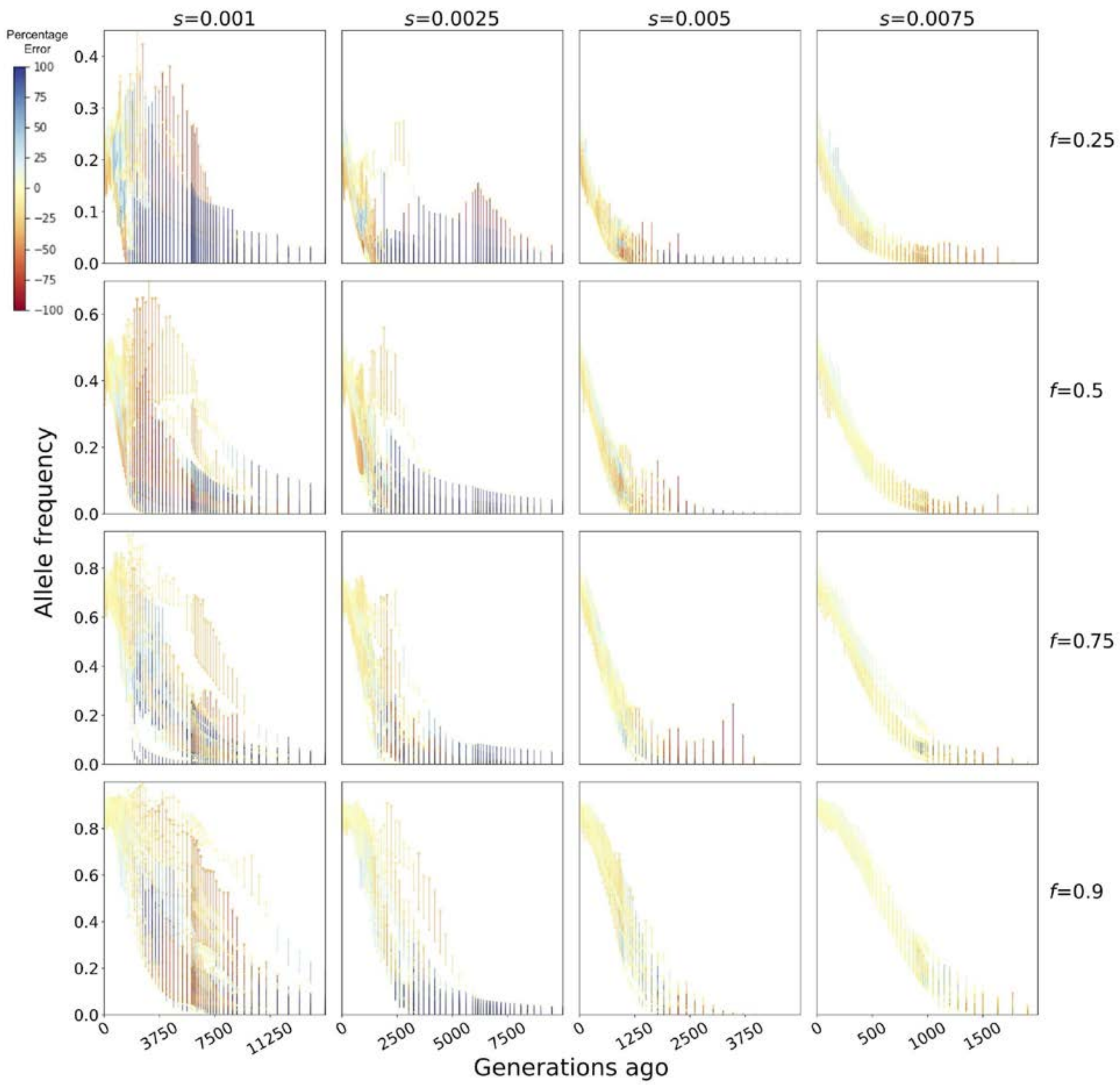


**Figure S7: Allele frequency (AF) trajectories inferred with SIA using true genealogies of simulations under the CEU demography.** Each panel shows 20 randomly selected examples of AF trajectories for a particular combination of selection coefficient and current AF. For each example, the true and inferred AF at each time point are connected by a vertical line with color scaled to the percentage error.

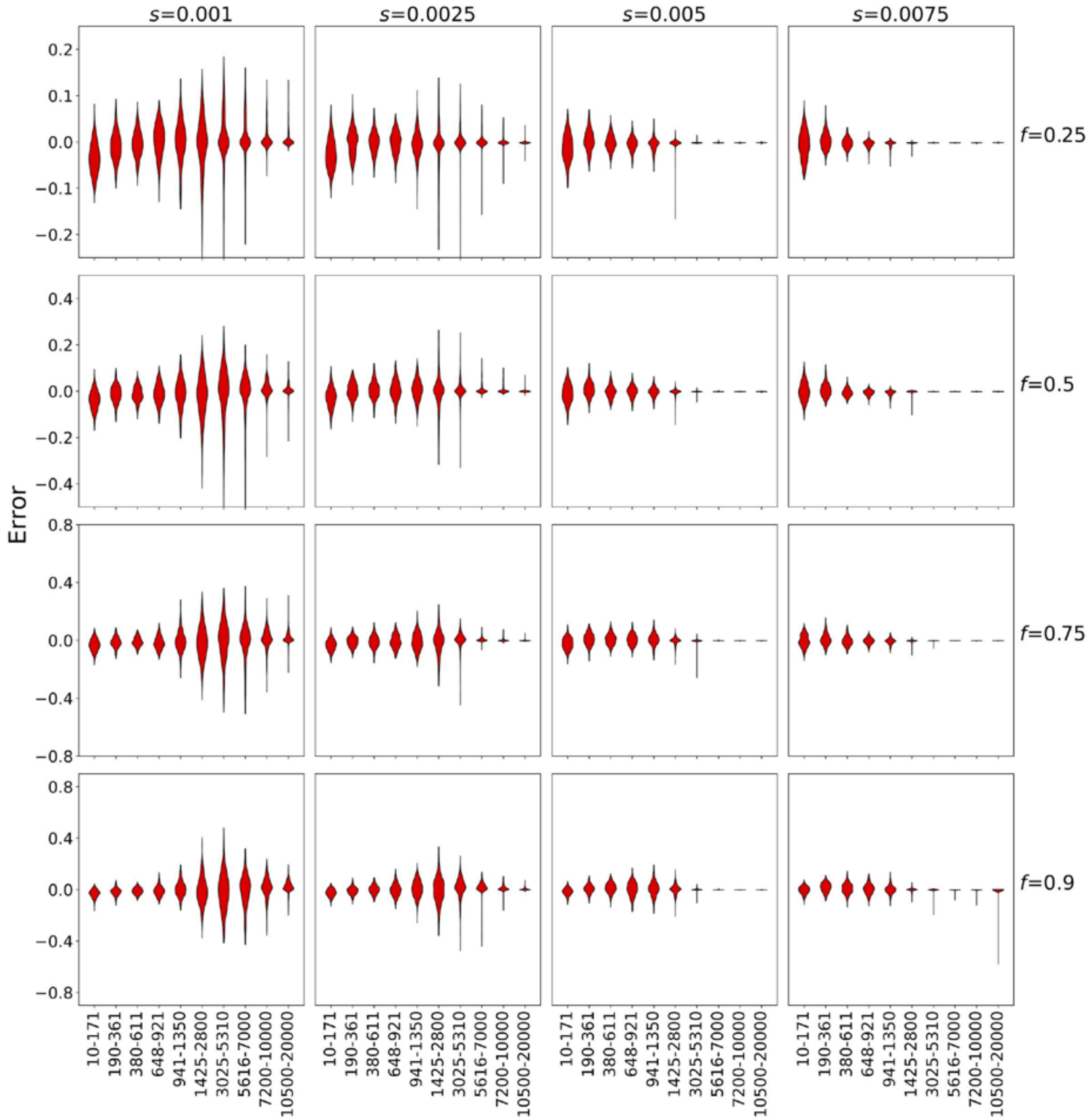


**Figure S8: Distribution of error in allele frequency trajectory inference with true genealogies of simulations under the CEU demography.** The performance of SIA for AF trajectory inference was evaluated with 100 simulations under each combination of selection coefficient and current AF. Violin plots in each panel show the distribution of absolute error in AF estimation binned for every 10 consecutive time points. Note that the *y*-axes limits for each row of panels with current AF equals *f* were set to be (-*f*, +*f*).

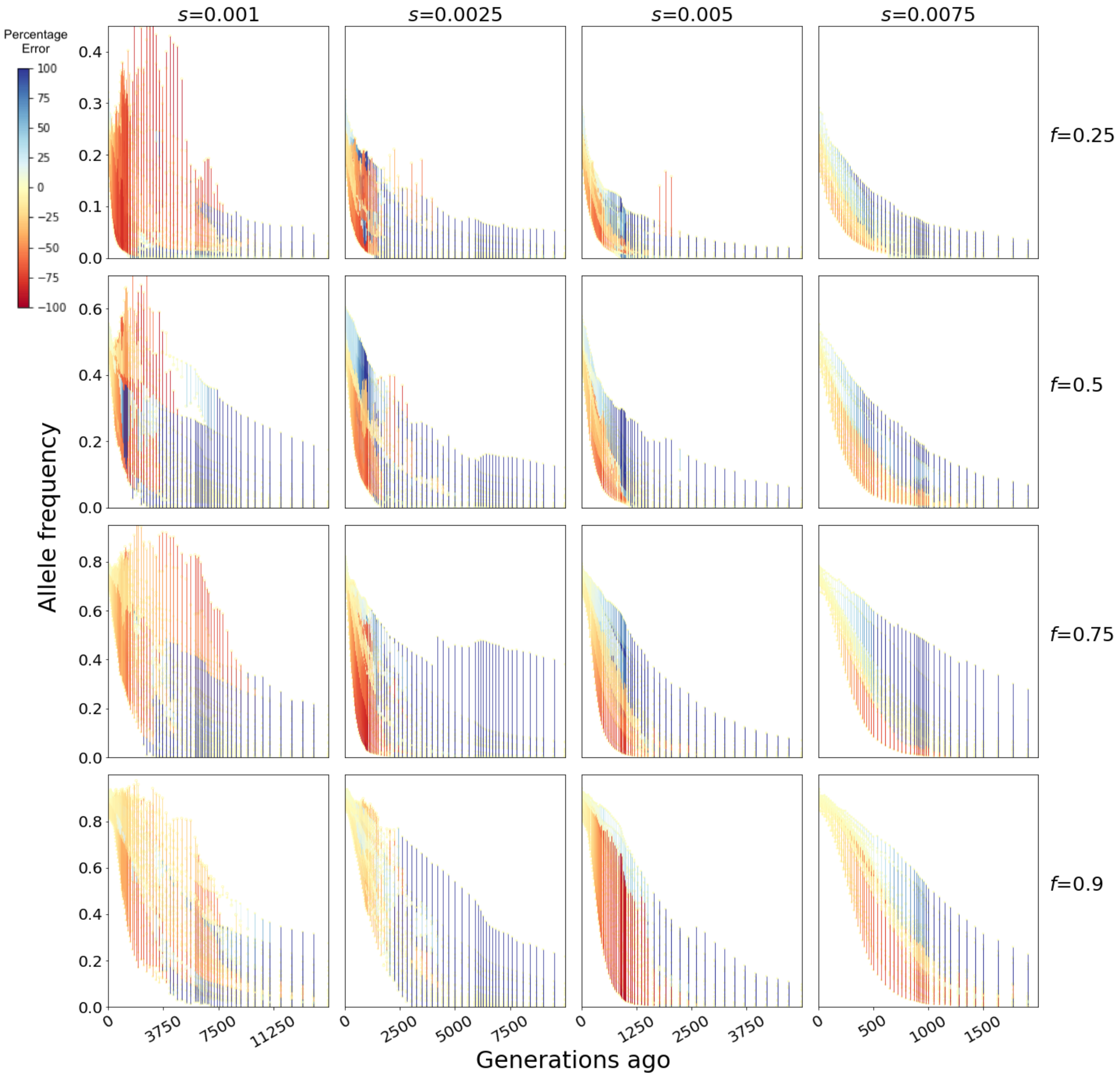


**Figure S9: Examples of allele frequency (AF) trajectories inferred with SIA using genealogies inferred from data simulated under the CEU demography.** Figure layout and description are identical to that of **Figure S7**.

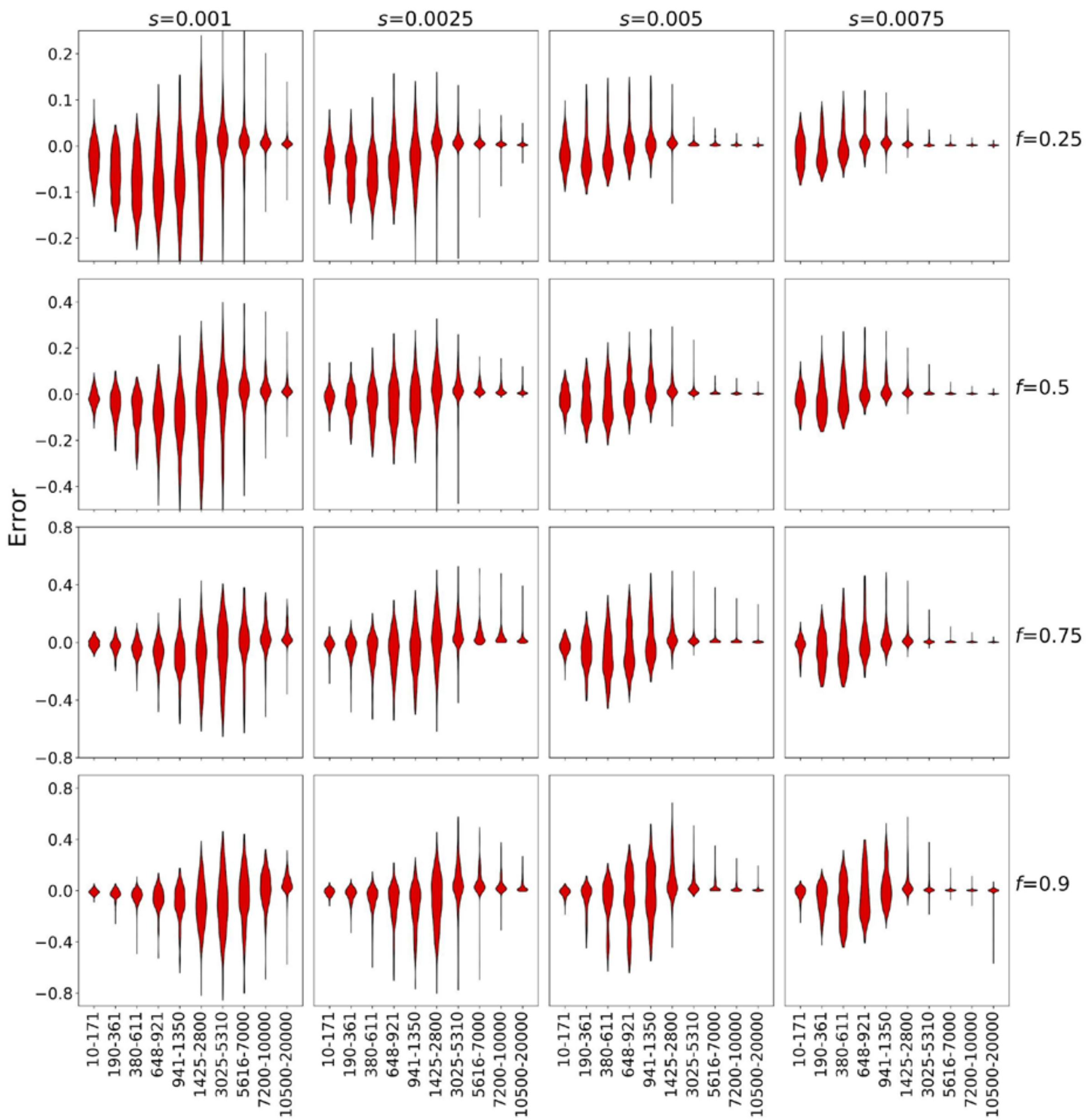


**Figure S10: Distribution of error in allele frequency trajectory inference with genealogies inferred from data simulated under the CEU demography.** Figure layout and description are identical to that of **Figure S8**.

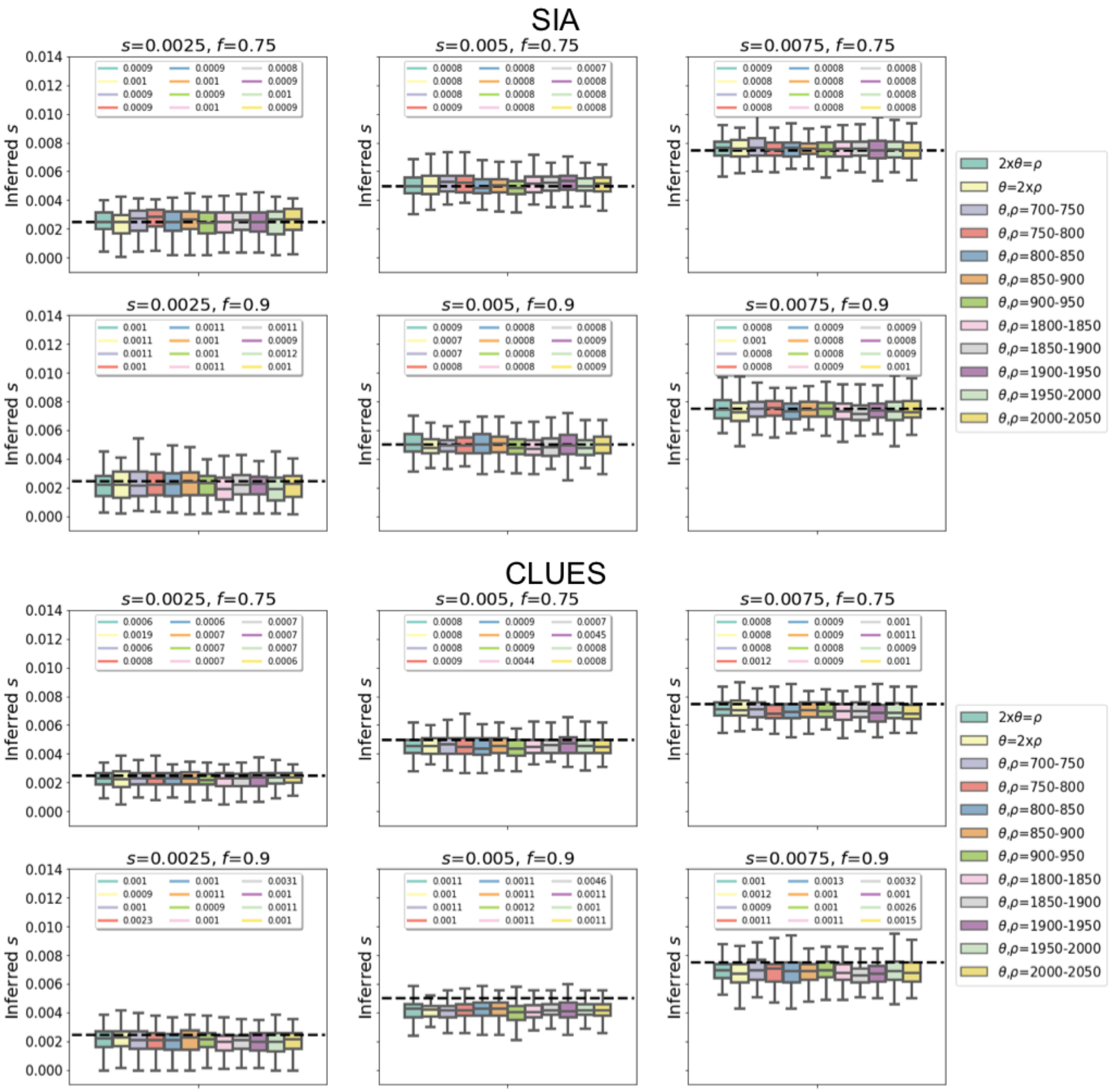


**Figure S11: Performance of SIA and CLUES models on selection coefficient inference, tested on true genealogies simulated under variable combinations of population-scaled mutation rate ($\theta$=$4N_e\mu L$) and population-scaled recombination rate ($\rho$=$4N_e rL$).** Each panel shows the model predictions for simulations of a particular selection coefficient (*s*) and current derived allele frequency (*f*). Each box represents a group of 100 simulations under $\theta$ and $\rho$ either specified by a fixed ratio, or sampled independently and uniformly within a particular range, as indicated in the legend. The dashed line marks the target value of *s*. The root mean squared error (RMSE) of the model predictions for each group of simulations is indicated at the top of the panel. For reference, the SIA model tested here was trained with true genealogies simulated under combinations of $\theta$ and $\rho$ sampled independently and uniformly from a range of [940, 1880].

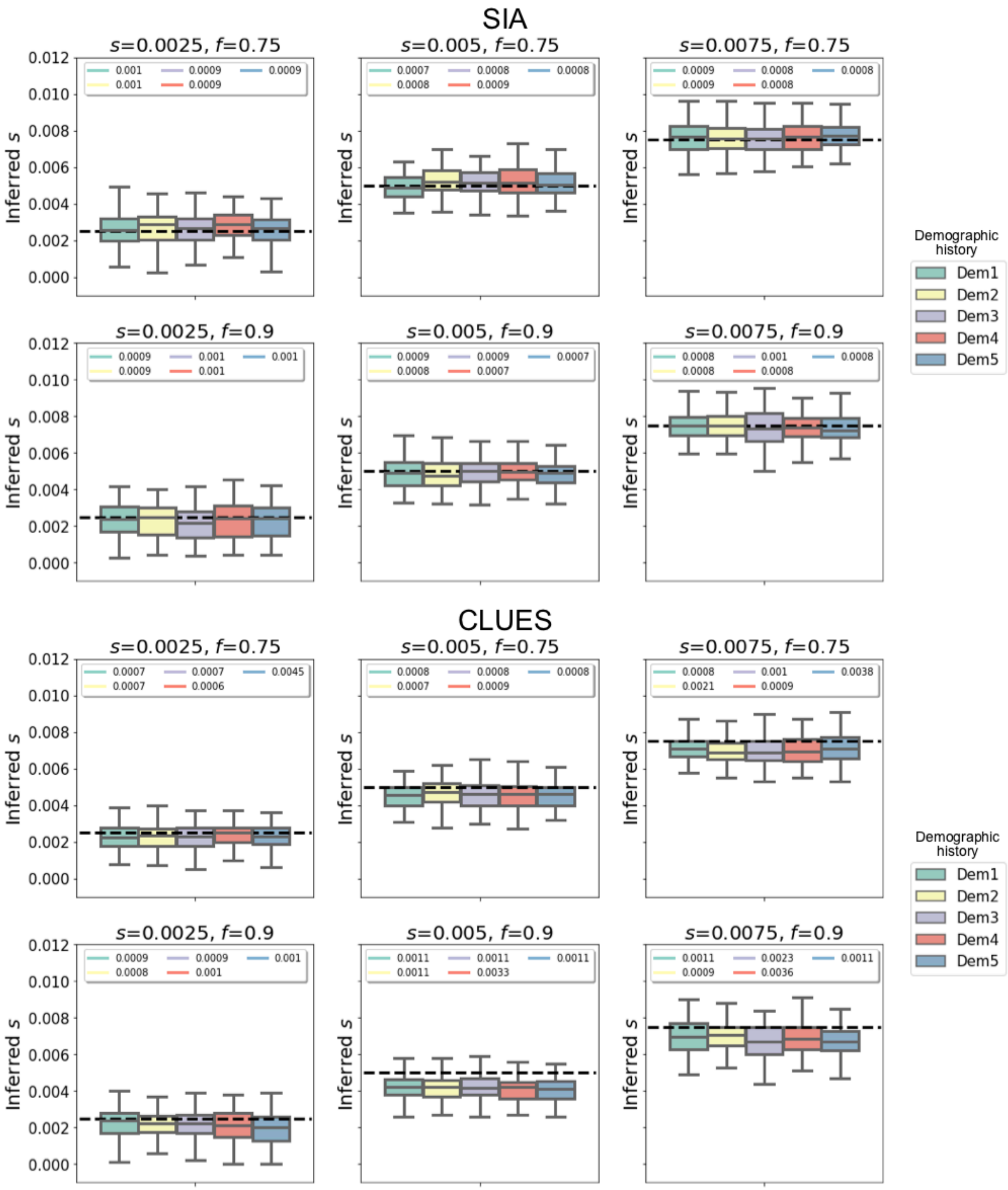


**Figure S12: Performance of SIA and CLUES models on selection coefficient inference, tested on true genealogies simulated under five alternative demographies.** Each demography is obtained from the CEU demography by modifying the population size at one time point during the recent population expansion phase (see **Figure S17** for more details). Each box represents a group of 100 simulations under the demography as indicated in the legend. Figure layout and description are otherwise similar to **Figure S11**.

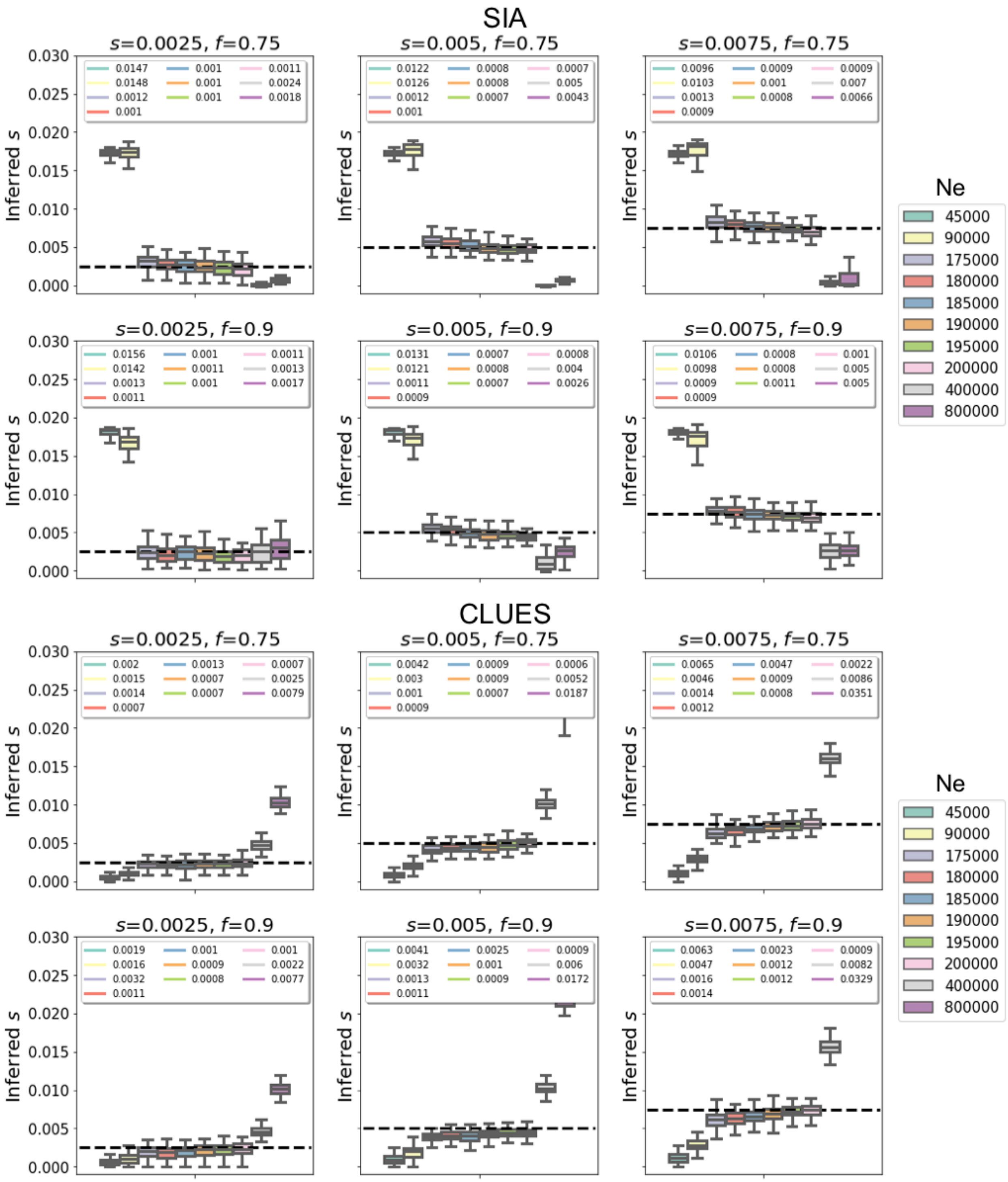


**Figure S13: Performance of SIA and CLUES models on selection coefficient inference, tested on true genealogies simulated under the CEU demography scaled to different present-day $N_e$.** Each group of 100 simulations of specific *s* and *f* under a particular present-day $N_e$ (i.e. a box) were performed with a globally scaled CEU demography such that the resulting demography has a present-day $N_e$ indicated by the legend (i.e. relative population size changes are preserved). For reference, the SIA model was trained with true genealogies simulated under a present-day $N_e$ of 188,088 (standard CEU). Figure layout and description are otherwise similar to **Figure S11**.

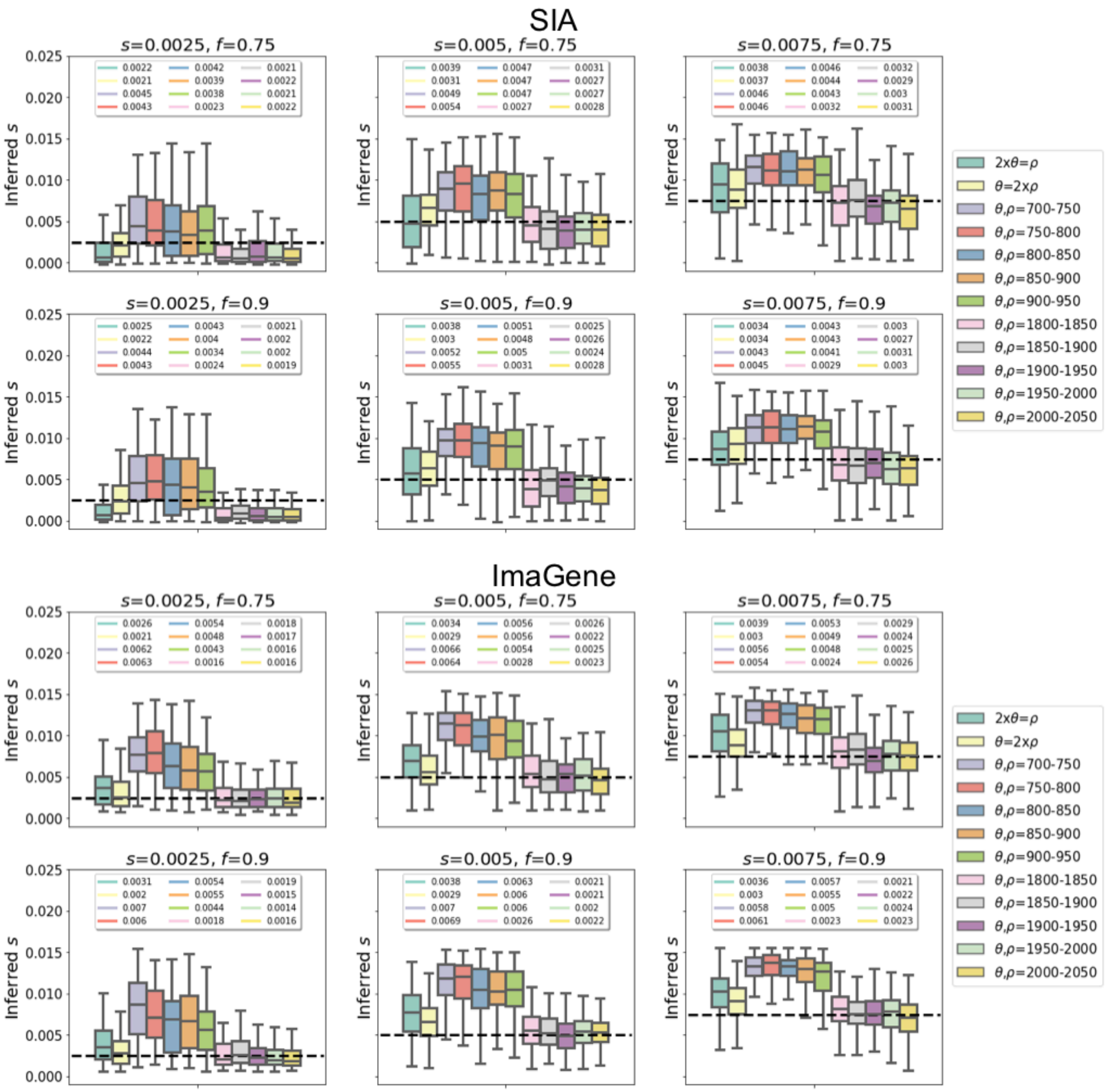


**Figure S14: Performance of SIA and ImaGene models on selection coefficient inference, tested on genealogies inferred by Relate from simulations under variable combinations of population-scaled mutation rate $\theta$ and population-scaled recombination rate $\rho$.** Note that the training data for both SIA and ImaGene are generated from identical sets of simulations, which were performed under combinations of $\theta$ and $\rho$ sampled independently and uniformly from a range of [940, 1880]. Figure layout and description are otherwise similar to **Figure S11**.

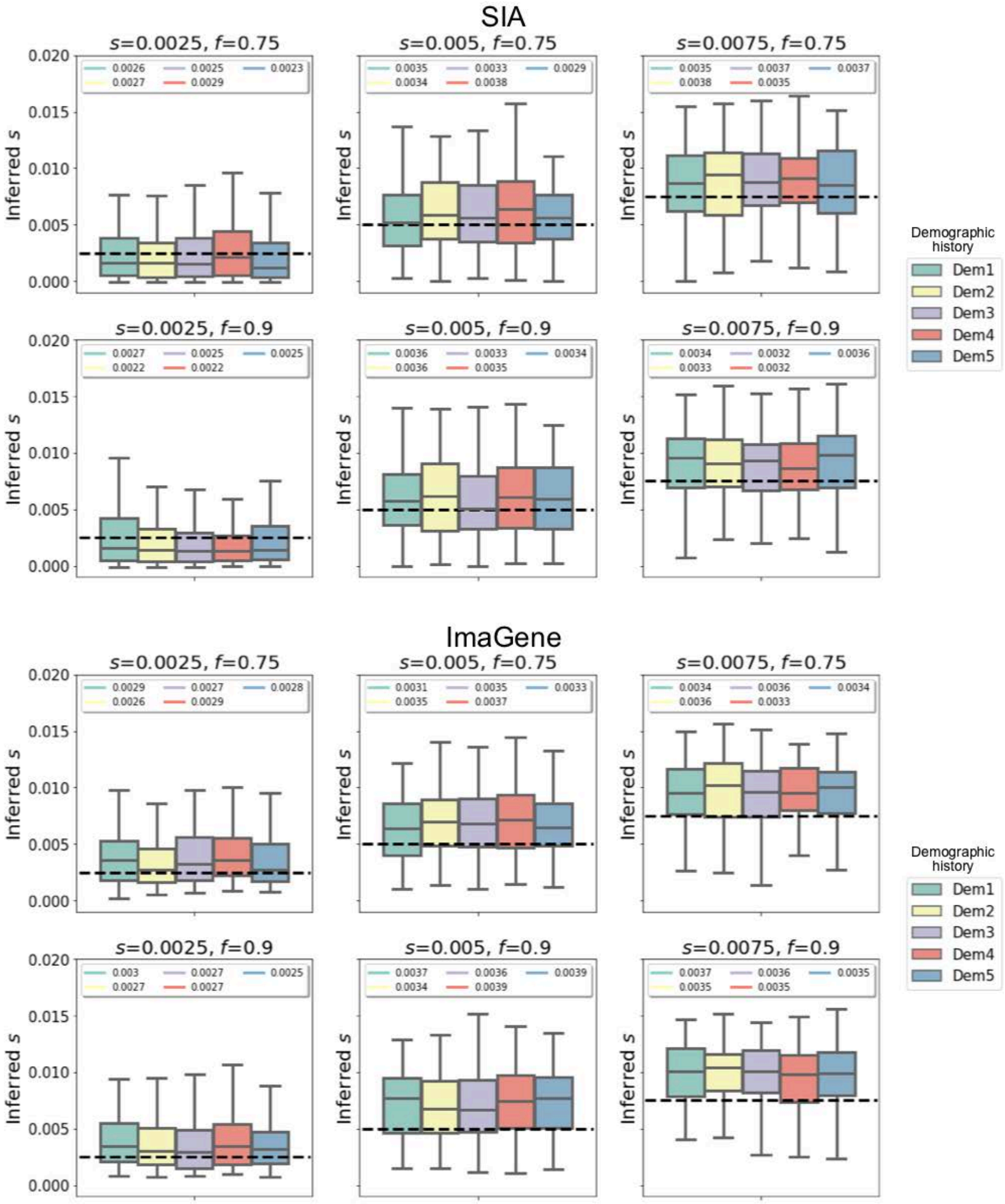


**Figure S15: Performance of SIA and ImaGene models on selection coefficient inference, tested on genealogies inferred by Relate from simulations under five alternative demographies.** Figure layout and description are otherwise similar to **Figure S12**. See **Figure S17** for details of the demography.

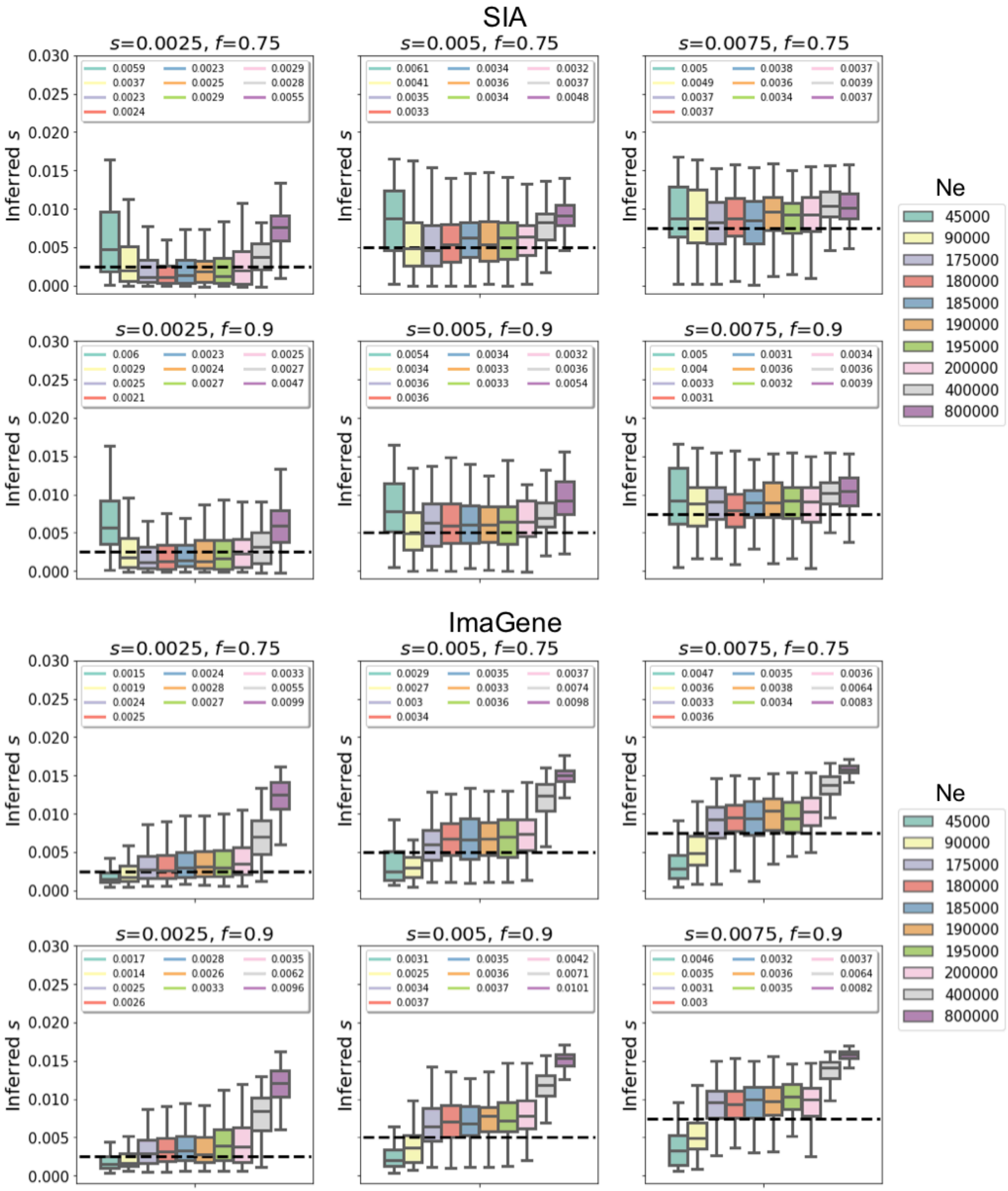


**Figure S16: Performance of SIA and ImaGene models on selection coefficient inference, tested on genealogies inferred by Relate from simulations under the CEU demography scaled to different present-day $N_e$.** Note that the training data for both SIA and ImaGene are generated from identical sets of simulations, which were performed under a present-day $N_e$ of 188,088 (standard CEU). Figure layout and description are otherwise similar to **Figure S13.**

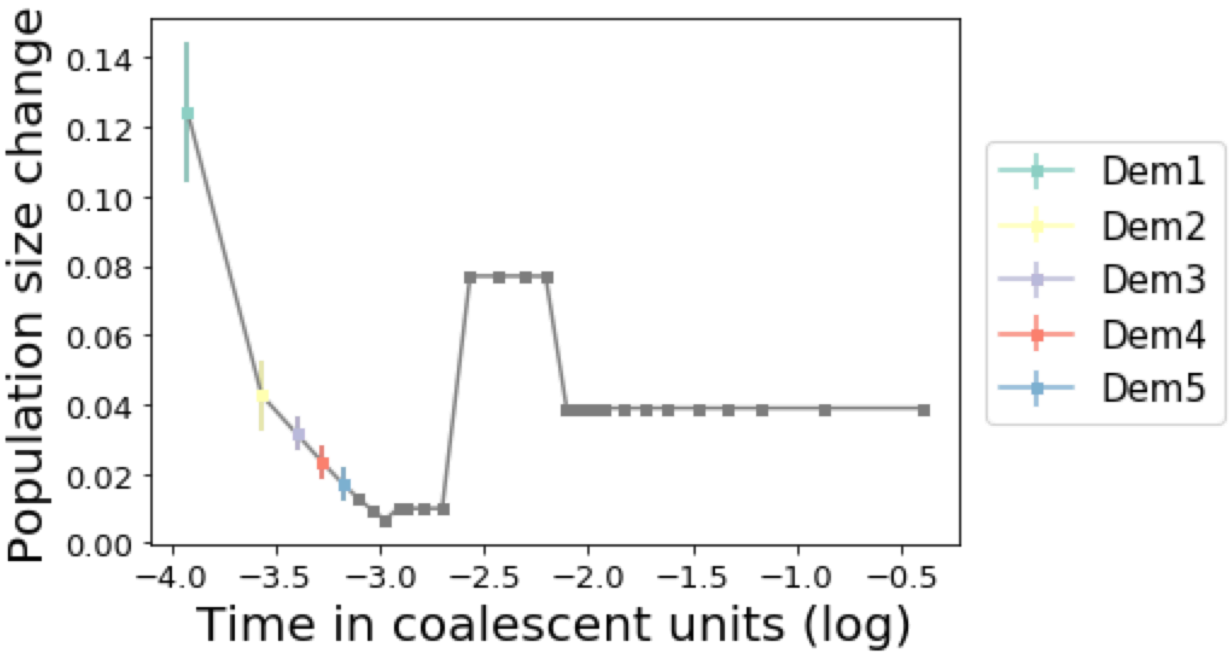


**Figure S17: Illustration of alternative demographies used to simulate test data plotted in Figure S12 and S15.** Squares indicate population size changes of the Tennessen *et al.* CEU model. For a simulation under a particular alternative demography, population size at the time point with matching color to the legend was modified by randomly sampling from a range centered on the original value ([-0.02, +0.02] for Dem1, [-0.01, +0.01] for Dem2, and [-0.005, +0.005] for Dem3-Dem5, as indicated by the vertical bar in the plot). Population sizes at all other time points were kept identical to the CEU demography.

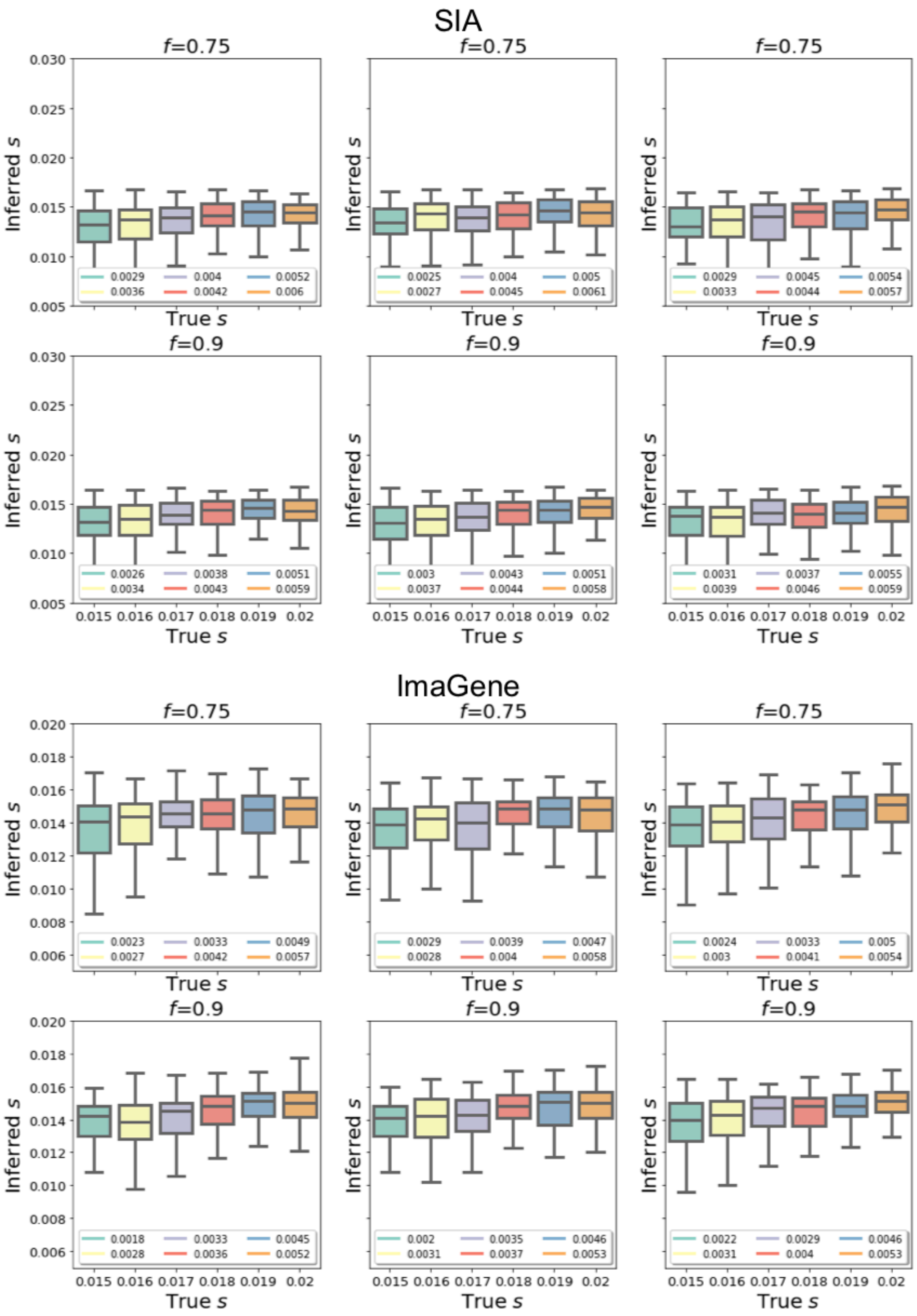


**Figure S18: Performance of SIA and ImaGene models on selection coefficient inference, tested on genealogies inferred by Relate from simulations under selection coefficients (*s*) beyond the range used for simulating training data.** Note that the training data for both SIA and ImaGene are generated from identical sets of simulations. Selection coefficients of sweep simulations constituting the training set were sampled uniformly from a range of [0.001, 0.02]. Figure layout and description are otherwise similar to **Figure S11**.

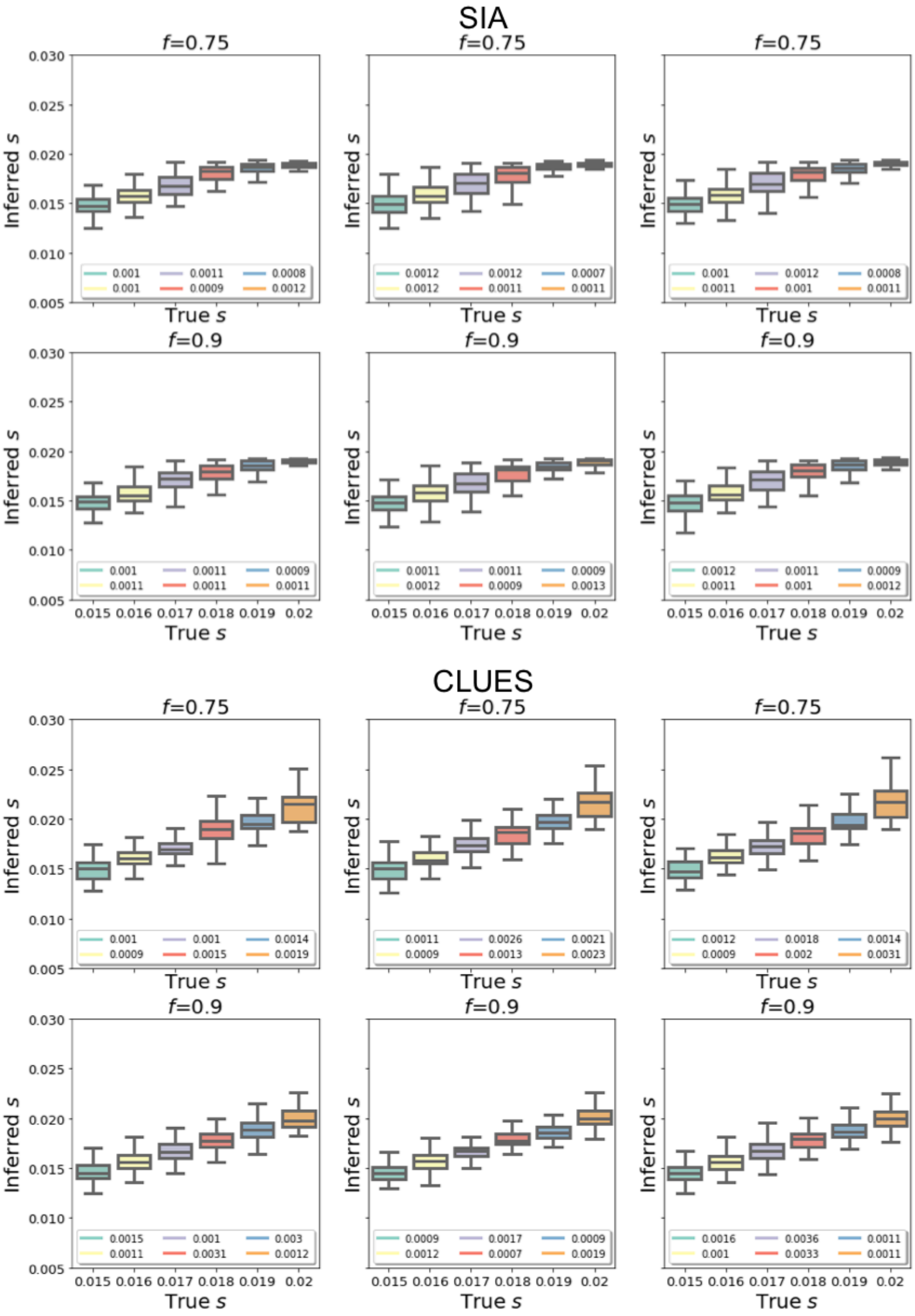


**Figure S19: Performance of SIA and CLUES models on selection coefficient inference, tested on true genealogies simulated under selection coefficients (*s*) beyond the range used for simulating SIA training data.** For reference, selection coefficients of sweep simulations constituting the SIA training set were sampled uniformly from a range of [0.001, 0.02]. Figure layout and description are otherwise similar to **Figure S18**.

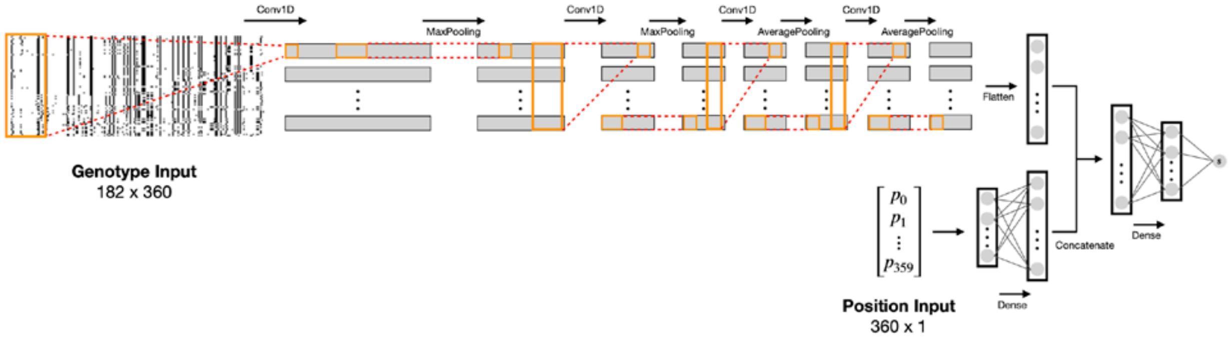


**Figure S20: Architecture of genotyped-based CNN for selection inference.** The model shares the exact same architecture as presented by Flagel et al. [14] with one modification — the original SoftMax output layer for classification was replaced by a linear output layer for selection coefficient inference.

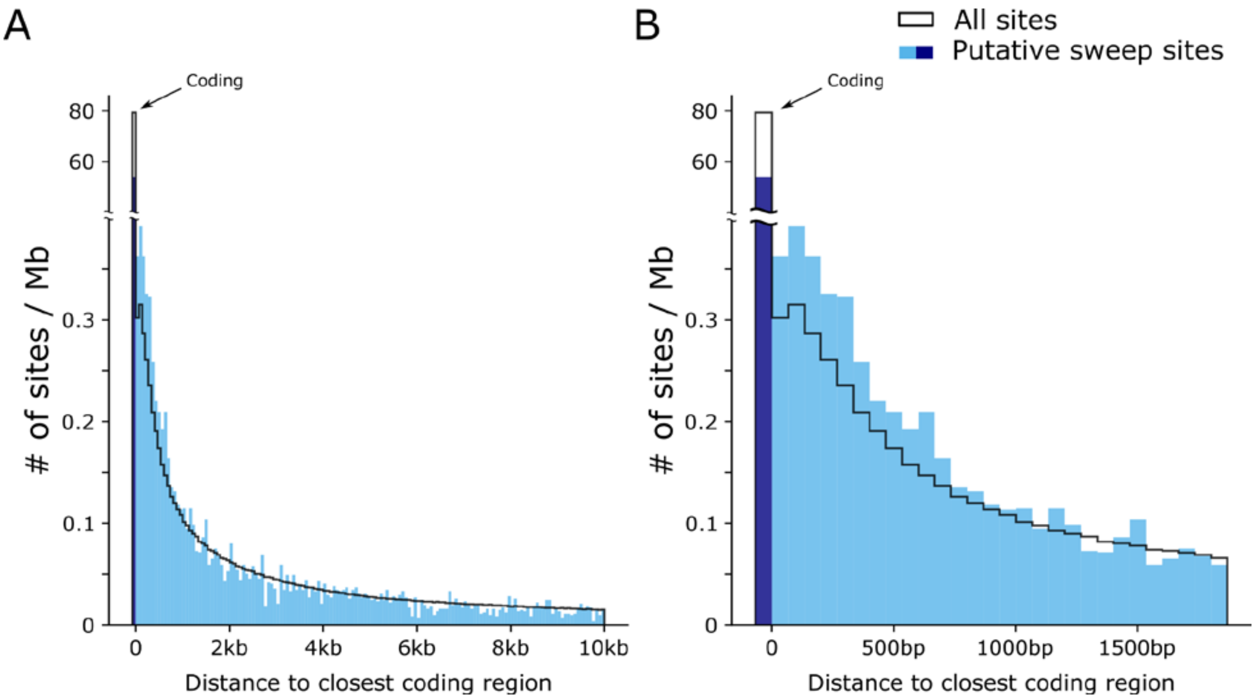


**Figure S21: Distribution of putative soft sweep sites in *S. hypoxantha* with respect to the nearest coding regions.** Panels (A) and (B) show the distribution of 15,551 sites (blue) across 333 scaffolds at different scales. For reference, the expected distribution of sites randomly drawn from all polymorphic sites is shown in black. Note that the sites that fall in coding regions are plotted in a separate bin.
